# Pericellular Matrix Proteoglycan Content is Less Affected in The Mid-Zone Cartilage in Early Post-Traumatic Osteoarthritis

**DOI:** 10.1101/2025.09.24.677987

**Authors:** Rasmus K. Hiltunen, Nina Pesonen, Simo P. Ojanen, Mikko A.J. Finnilä, Sanna Palosaari, Walter Herzog, Simo Saarakkala, Rami K. Korhonen, Juuso T. J. Honkanen, Petri Tanska

## Abstract

Proteoglycan content changes in articular cartilage during post-traumatic osteoarthritis affect both chondrocyte biomechanics and tissue health. Previously, we found that four weeks after anterior cruciate ligament transection (ACLT) in rabbits, proteoglycan loss was less pronounced in the pericellular matrix (PCM) than in the extracellular matrix (ECM), resulting in a greater PCM-to-ECM proteoglycan content ratio compared to controls. Here, we investigate whether this pattern is already present two weeks post-ACLT and whether changes in the proteoglycan ratio relate to cell volumetric strain in the superficial zone during tissue loading.

Unilateral ACLT was performed in eight skeletally mature rabbits. Two weeks post-surgery, cartilage from all load-bearing sites of the operated and contralateral knees were collected. Age-matched, non-operated knees from separate animals served as controls. Proteoglycan content in the chondrocyte microenvironment was analyzed in the superficial and middle zones of cartilage via digital densitometry. Volumetric strain of superficial zone cells was measured using two-photon confocal microscopy during indentation.

In the superficial zone, the PCM-to-ECM proteoglycan content ratio was greater in the ACLT knees than in controls for the lateral tibial plateau (*p* = 0.043). However, this was not clearly linked to changes in cell volumetric strain. Interestingly, in the middle zone, the PCM-to-ECM proteoglycan content ratio was greater in the ACLT knees than in controls for the medial femoral condyle, femoral groove, and patella (*p* < 0.05). These results suggest that, in early post-traumatic osteoarthritis, proteoglycan loss is less severe in the PCM than in the ECM, especially in the middle cartilage zone.

## 1. Introduction

Chondrocytes (cartilage cells) maintain and repair the extracellular matrix (ECM) of articular cartilage by converting external physical stimuli into biochemical responses. This mechanism is known as mechanotransduction [1]. The ECM primarily consists of a fibrous collagen network, proteoglycans, interstitial fluid, and non-collagenous proteins [2]. The highly organized collagen network provides tensile strength to the tissue, while proteoglycans in the interfibrillar space contribute significantly to compressive stiffness [3].

Osteoarthritis (OA) is the most common knee joint disease, and it significantly impacts patients’ quality of life [4]. OA disrupts cartilage homeostasis, favoring catabolic processes [5]. Abnormal cartilage loading is a key contributor to OA development [6], triggering chondrocyte dysfunction and apoptosis [7]. Both excessive and insufficient cartilage strains can elevate the production of inflammatory and catabolic mediators, which both suppress matrix synthesis and promote loss of proteoglycans and collagen [8]. Interestingly, early OA may also involve a localized healing response, enhancing proteoglycan production [9,10] and potentially slowing down or even partially reversing matrix degradation [11,12]. These local biomechanical and biochemical changes may influence how chondrocytes perceive and respond to mechanical signals [13,14]. Therefore, understanding how the tissue matrix is altered and how those alterations contribute to cell strains and responses in very early stages of OA is crucial to unraveling the disease pathogenesis.

Chondrocytes are surrounded by the pericellular matrix (PCM), a specialized microdomain rich in collagen type VI and specific proteoglycans such as perlecan and decorin [14]. Together with the territorial ECM, they form a distinct microenvironment [15,16]. Due to its proximity to chondrocytes, the PCM is thought to mediate both biomechanical and biochemical signals [17,18]. Computational and experimental studies have suggested that proteoglycans in the PCM influence chondrocyte deformations and mechanotransduction [19–22]. Although experimental data are limited, findings from animal models [23] and human studies [24] suggest that changes in pericellular proteoglycan and collagen content may precede alterations in the broader ECM.

Animal models help study OA pathogenesis in a controlled setting, minimizing variability found in human studies (e.g., age, gender, and weight) [25]. In rabbits, anterior cruciate ligament transection (ACLT) is commonly used to induce knee joint instability, quickly and reliably producing post-traumatic OA with cartilage degradation patterns similar to those observed in humans [26]. In rabbits, site-specific changes in the superficial (and middle) zone ECM proteoglycan content can be detected as early as four weeks post-ACLT, accompanied by disorganization of the collagen network in the superficial and middle zone cartilage [27,28]. Within the chondrocyte microenvironment, the proteoglycan content of the PCM degraded less than that of the surrounding ECM, indicated as a greater PCM-to-ECM proteoglycan content ratio in the operated knees compared to non-operated controls [12]. More recently, we also reported that ECM proteoglycan loss is detectable already two weeks post-ACLT while the collagen network remained largely intact [29]. Moreover, we detected changes in the superficial zone cell shape and volume change following mechanical loading of the tissue in ACLT-operated knees compared to non-operated control knees that we could not explain with the proteoglycan content changes in the ECM.

Thus, our primary aim was to determine whether changes in the proteoglycan content within the chondrocyte microenvironment at two weeks post-ACLT cartilage resemble those observed at four weeks post-ACLT. Specifically, we examined whether proteoglycan loss in the PCM of the superficial and middle cartilage zones is less than in the surrounding ECM, as indicated by a greater PCM-to-ECM content ratio. In a secondary analysis, we compared superficial zone pericellular proteoglycan content with previously reported chondrocyte shape and volume change data from the same sample set [29] to explore a potential link between the pericellular proteoglycan content, and cell morphology and volume change following mechanical loading of the tissue.

## 2. Methods

### 2.1. Animal model

A unilateral ACLT surgery was performed in eight skeletally mature female New Zealand white rabbits (*Oryctolagus cuniculus*, age 12 months at the time of surgery). The operated knee joint was chosen randomly to avoid bias [29]. The rabbits were euthanized two weeks post-surgery. Both operated (ACLT) and contralateral (CL) knee joints were harvested. Randomly selected knee joints (left or right leg) from eight age-matched and unoperated female rabbits were collected for the control group (CNTRL). Osteochondral samples were prepared from lateral and medial femoral condyles, lateral and medial tibial plateaus, femoral groove, and patella (Figure 1A-D). All procedures were approved by the Committee on Animal Ethics at the University of Calgary and were carried out according to their guidelines (ethical permit #AC11-0035). After tissue harvest, a simultaneous two-photon confocal microscopy and *in-situ* indentation testing [30] was performed for the main load-bearing regions of cartilage under a force-relaxation protocol to evaluate chondrocyte shape and volume before and after loading. Following the *in-situ* indentation and confocal microscopy, reported in detail in our previous study [29] (see section 2.4 for a brief overview), samples were fixed in formalin, decalcified, dehydrated in an ascending alcohol series, embedded in paraffin, and processed for polarized light microscopy and digital densitometry.

**Figure 1.**
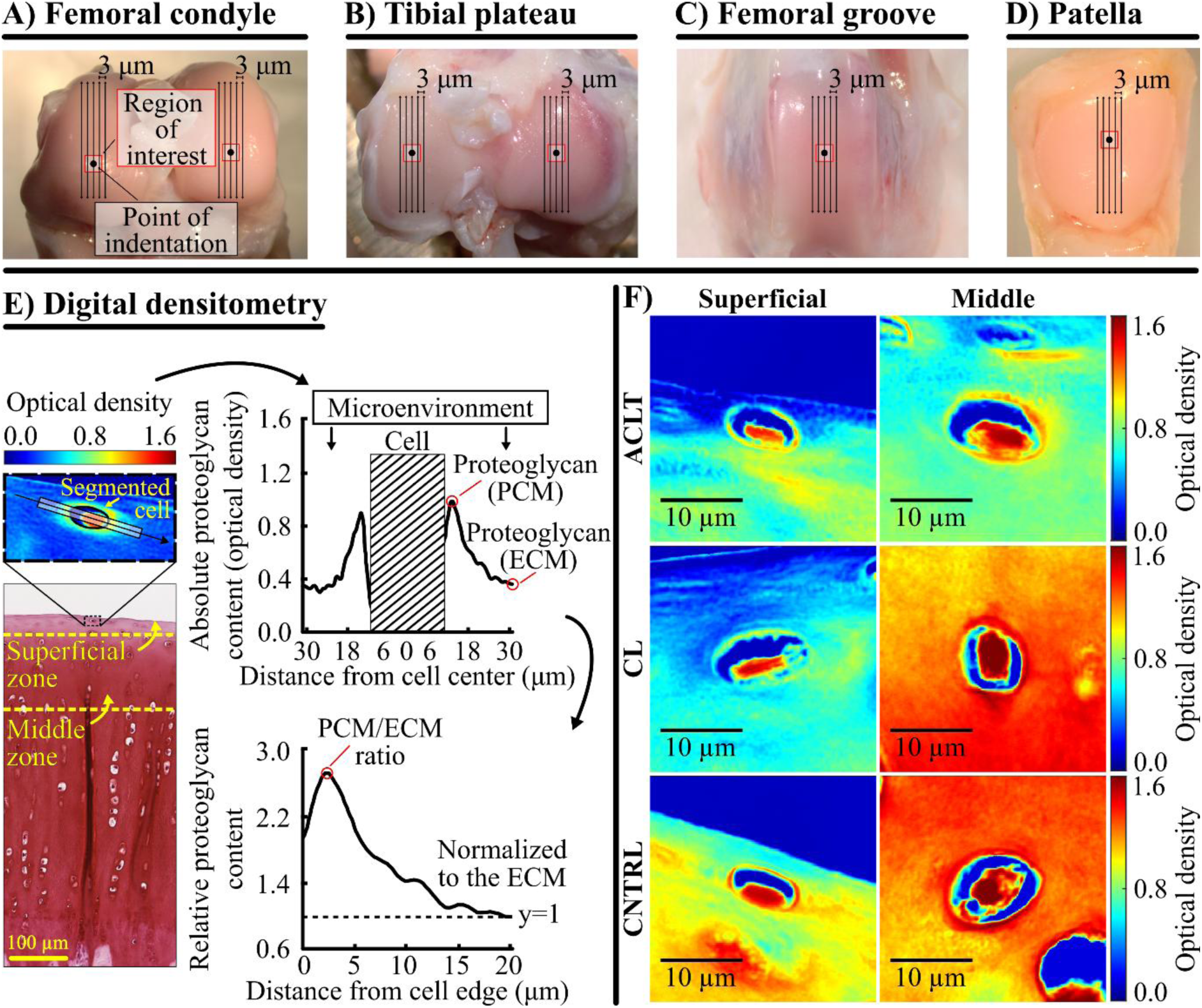
The knee joints of skeletally mature rabbits were harvested two weeks after anterior cruciate ligament transection surgery. Samples were prepared from A) lateral and medial femoral condyles, B) lateral and medial tibial plateaus, C) lateral femoral groove, and D) patella. The primary load-bearing areas of each cartilage surface were defined as regions of interest: the highest (proximal) point of femoral condyles and the centers of tibial plateaus, femoral groove, and patella. E) Histological sections were prepared as shown in subfigures A-D and digital densitometry was used to measure Safranin-O-stained histological sections to analyze the proteoglycan content (optical density, scaled from 0 to 3.0) of the cell microenvironment in both superficial and middle zone cartilage. Proteoglycan content was analyzed from a rectangular region of interest extending 20 µm from the cell edge (height = 6 µm). The peak proteoglycan content value within 5 µm of the cell edge was designated for the pericellular matrix (PCM). The value 20 μm from the cell edge was set to represent the extracellular matrix (ECM). The PCM/ECM ratio was calculated to highlight changes in the cell microenvironment relative to the ECM. F) Examples of optical density images acquired with digital densitometry from the femoral groove. ACLT, Anterior cruciate ligament transection; CL, contralateral; CNTRL, control.

### 2.2. Polarized light microscopy and digital densitometry

Three unstained sections (5 µm thick) per sample were prepared for polarized light microscopy from the areas where the *in-situ* indentation testing was performed. Polarized light microscopy was used to measure the average depth-wise collagen orientation angle profile for the CNTRL group rabbits to define the superficial and middle zones at each cartilage location [29]. Furthermore, three Safranin-O-stained sections (3 μm thick) per sample were prepared from the same locations. The stained sections were imaged first with light microscopy to select superficial and middle zone chondrocytes for digital densitometry (Figure 1F). Chondrocytes that were visibly intact and located at least 30 µm from the nearest adjacent chondrocyte were chosen from the same areas where the indentation test was performed. Then, the proteoglycan content of the cell microenvironment (PCM and surrounding ECM) was estimated via digital densitometry measuring the optical density of the Safranin-O-stained sections [12]. This method has been originally validated with biochemistry [31,32]. The number of cells imaged are shown in Supplementary Table 1. The measurement setup consisted of a light microscope (Microphot FXA, Nikon Co., Tokyo, Japan) connected to a CCD digital camera (ORCA-ER, Hamamatsu Photonics K.K., Hamamatsu, Japan). Monochromatic light (λ: 492 ± 5 nm) and 40x magnification objective (NA: 0.7) were used to give a spatial resolution of 0.43×0.43 μm^2^. After collection, the resulting grayscale images (1024×1344 pixels) were converted into optical density images using a calibration line obtained with optical density filters (optical density values of air, 0.0, 0.3, 0.6, 1.0, 1.3, 1.6, 2.0, 2.3, 2.6, 3.0) and defined for each pixel of the CCD camera.

### 2.3. Analysis of the chondrocyte microenvironment

Chondrocytes were segmented by overlaying the digital densitometry images onto the light microscopy images (Supplementary Figure 1). Absolute proteoglycan content profiles of the chondrocyte microenvironment were averaged from a rectangular region (height: 6 μm) that extended 20 μm from the cell border towards the ECM on both sides of the cell (Figure 1E). The region was aligned parallel to the cell’s major axis. The profiles were used to calculate the peak proteoglycan content value within 5 µm of each cell’s edge, a region that presumably contains the PCM [33]. Secondly, we calculated the PCM-to-ECM proteoglycan content ratio by dividing the peak proteoglycan content value by the value located at 20 μm from the cell’s edge (ECM). This normalization provides a quantitative value for the changes in the cell microenvironment relative to the ECM. Lastly, the location of the cell from the cartilage surface and the location of the peak proteoglycan content from the cell edge were analyzed (Supplementary Tables 2 and 3).

### 2.4. In-situ two-photon confocal microscopy and indentation testing

These measurements were previously performed in our earlier study [29]. Briefly, simultaneous two-photon confocal microscopy and *in-situ* indentation testing [30] were performed on the main load-bearing regions of the cartilage after tissue harvest to determine how chondrocyte volume response to loading is altered two weeks post-ACLT surgery (Figure 2). Prior to the microscopy tests, the ECM and dead cells were stained with fluorescently conjugated Dextran (D34682, Thermo Scientific, Eugene, Oregon, USA) and Propidium Iodide, respectively.

**Figure 2.**
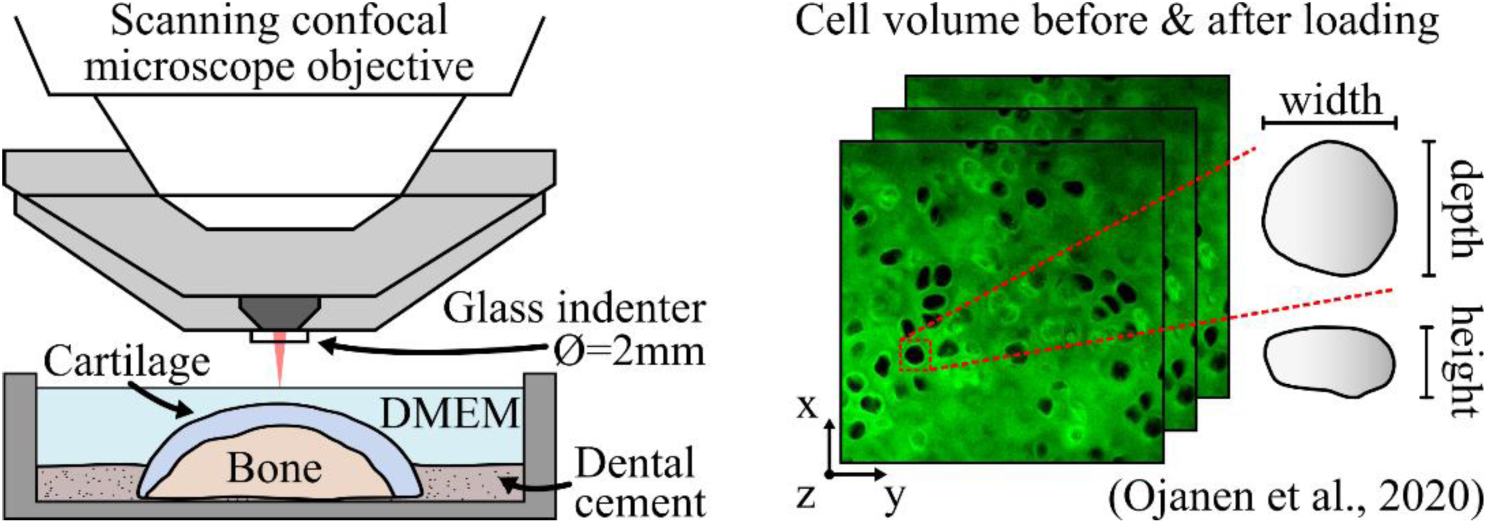
In our previous study [29], confocal microscopy imaging was performed on cartilage samples stained with Dextran (matrix) and Propidium iodide (dead cells) before and after force-relaxation loading in indentation. DMEM, Dulbecco’s Modified Eagle Medium.

First, a pre-load of 0.1 – 0.2 MPa was applied and maintained for 20 minutes, after which a “before loading” image stack was collected. The tissue was then indented at a rate of 10 µm/s until a load of 2 MPa was achieved. Subsequently, the indenter displacement was held constant for 20 minutes to allow full tissue relaxation, followed by acquisition of an “after loading” image stack at the same location. For each group and each location, approximately 40 – 70 cells were measured and analyzed using open source segmentation/analysis software pyCellAnalyst (http://pycellanalyst.readthedocs.io/en/latest/index.html). As measurements were performed on the intact cartilage surface, the cell volume analysis covered only the superficial zone of cartilage (depth penetration is limited to ∼60–100 µm depending on the excitation wavelength, fluorophore, and laser power). Cell volumetric strain caused by the mechanical loading was analyzed as *ε*_vol_ = (*V*_after_ − *V*_before_)/ *V*_before_, where *V_before_* and *V_after_* are the volumes measured before and after loading, respectively. Further details and complete results are presented in the Supplementary material, section A, and in Ojanen et al. [29].

### 2.5. Statistical analysis

A linear mixed-effects (LME) model was used to compare the PCM-to-ECM proteoglycan content ratio and the peak proteoglycan content (located in the PCM) between the experimental groups. In the LME model, the experimental group, normalized depth of the cell, and side of the profile (“left” or “right”) were set as fixed effects. The samples were treated as a random effect to consider the within-animal dependence among the operated rabbits. For pairwise comparisons, the least significant difference was used. Unless stated otherwise, results are presented as means ± 95% confidence intervals from the LME model. Statistical analyses were performed using IBM Statistics (version 29, IBM, Armonk, NY, USA) and MATLAB (R2023a, MathWorks Inc., Natick, MA, USA), with the level of significance set to α = 0.05.

## 3. Results

### 3.1 Pericellular matrix to extracellular matrix proteoglycan content ratio

The PCM-to-ECM proteoglycan content ratio of the ACLT group knees was 15.0% [CI95: 0.5, 29.5%, *p* = 0.043] greater compared to the CNTRL group knees for the superficial zone cartilage of the lateral tibial plateau (Figure 3A.4). The PCM-to-ECM ratio in the lateral femoral condyle of the ACLT group knees was 20.1% [CI95: 2.1, 38.1%, *p* = 0.033] greater than that of the CL group knees. In addition, the PCM-to-ECM ratio of the CL group knees was 26.3% [CI95: 4.7, 47.8%, *p* = 0.020] greater compared to the CNTRL group knees.

**Figure 3.**
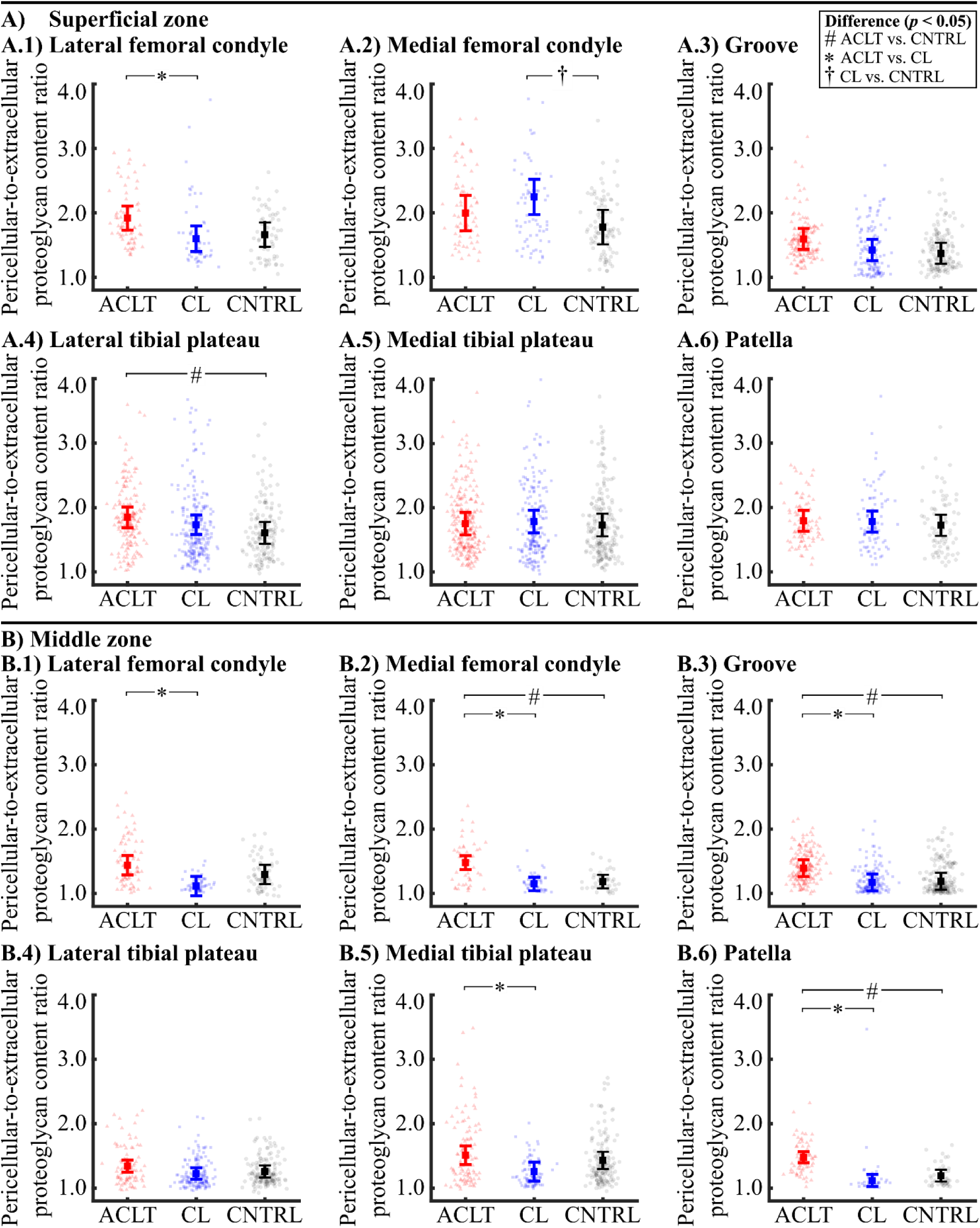
Pericellular-to-extracellular proteoglycan (PCM-to-ECM) content ratio of the A) superficial and B) middle zone cell microenvironment in the 1) lateral femoral condyle, 2) medial femoral condyle, 3) lateral femoral groove, 4) lateral tibial plateau, 5) medial tibial plateau, and 6) patella. The mean and 95 % confidence intervals for each group are shown. Black points indicate the individual values. ACLT, anterior cruciate ligament transection; CL, contralateral; CNTRL, control.

In the middle zone cartilage, differences in the PCM-to-ECM proteoglycan content ratio between the ACLT and CNTRL group knees were more pronounced than in the superficial zone (Figure 3B). The PCM-to-ECM ratio of the ACLT group knees was greater than in the CNTRL group knees for the medial femoral condyle (24.8% [CI95: 11.9, 37.6%], *p* < 0.001), the femoral groove (17.1% [CI95: 1.6, 32.6%], *p* = 0.033), and the patella (24.0%, [CI95: 13.5, 34.4%], *p* < 0.001). Additionally, the PCM-to-ECM ratio was 19.0-32.4% greater in the ACLT group knees than in the CL group knees at each knee location except for the lateral tibial plateau.

### 3.2. Peak proteoglycan content of the cell microenvironment

The peak proteoglycan content (located in the PCM) was 34.6% [CI95: −58.5, −10.8%, *p* = 0.007] and 32.6% [CI95: −53.7, −11.4%, *p* = 0.005] smaller in the superficial zone cartilage of the lateral tibial plateau and patella of the ACLT group compared to the CNTRL group (Figure 4A). Also, the peak proteoglycan content was 33.9–42.9% smaller in the lateral femoral condyle, [−39.5%, CI95: −65.3, −13.7%, *p* = 0.008], the lateral tibial plateau [−34.0%, CI95: - 50.6, −17.4%, *p* = 0.002], the medial tibial plateau [−33.9%, CI95: −57.4, −10.3%, p = 0.007], and the patella [−42.9%, CI95: −52.3, −33.5%, *p* < 0.001] in the ACLT group compared to the CL group rabbits. The peak proteoglycan content was the same for the CL and CNTRL group rabbits for all locations of the superficial zone cartilage.

**Figure 4.**
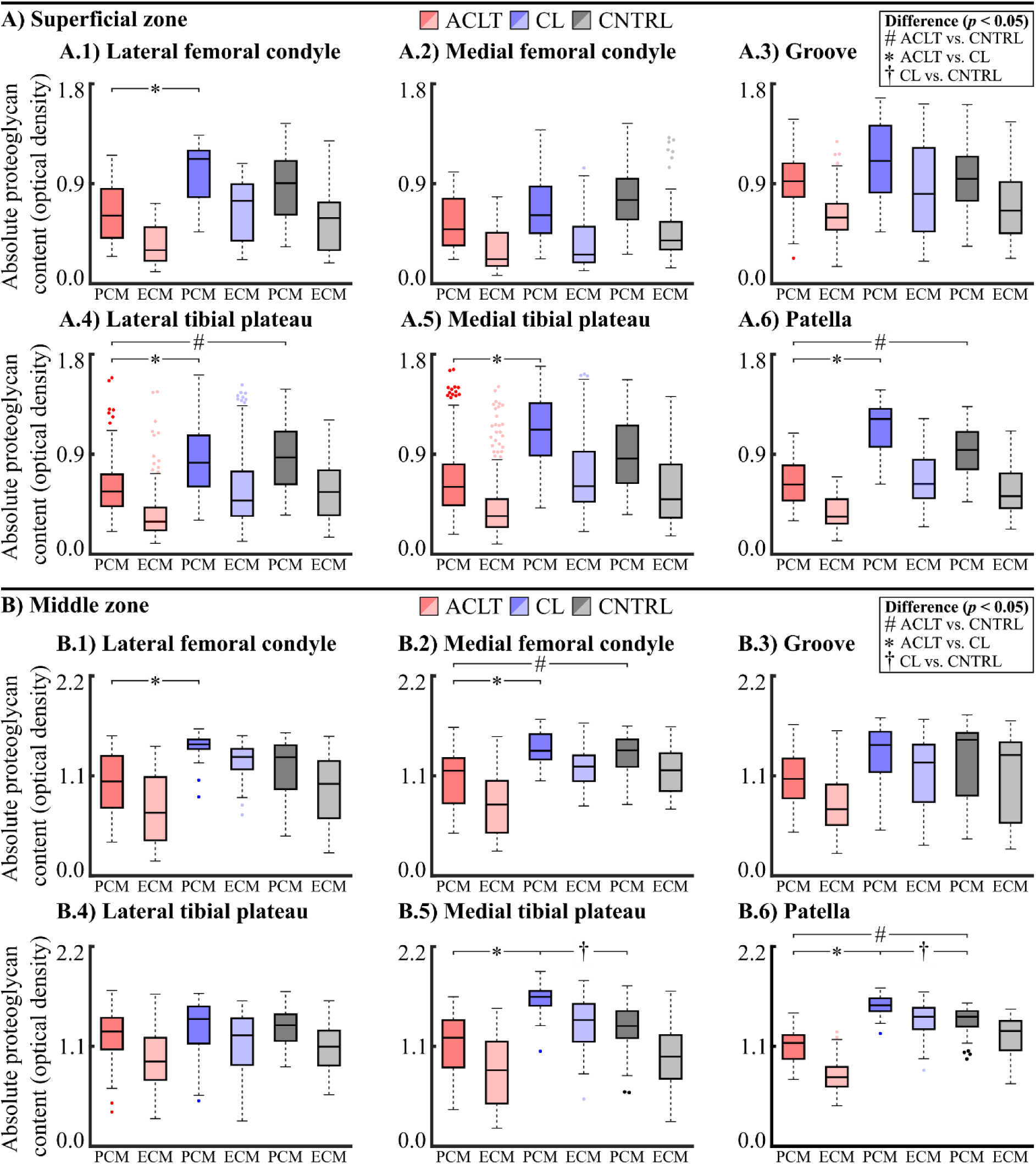
Box and whiskers plot for peak (PCM, darker-colored boxes) and extracellular (ECM, dim-colored boxes) proteoglycan content of the A) superficial and B) middle zone cell microenvironment in the 1) lateral femoral condyle, 2) medial femoral condyle, 3) lateral femoral groove, 4) lateral tibial plateau, 5) medial tibial plateau, and 6) patella. Circles represent outliers. ACLT, anterior cruciate ligament transection; CL, contralateral; CNTRL, control.

The peak proteoglycan content was 22.3% [CI95: −36.1, −8.4%, *p* = 0.003] and 20.6% [CI95: - 30.1, −11.1%, *p* < 0.001] smaller for the middle zone cartilage of the medial femoral condyle and patella of the ACLT compared to the CNTRL group rabbits (Figure 4B). Also, the peak proteoglycan content of the ACLT group rabbits was 23.8-29.8% smaller than that of the CL group rabbits for the lateral femoral condyle [−27.6%, CI95: −45.6, −9.7%, *p* = 0.008], the medial femoral condyle [−24.4%, CI95: −35.2, −13.6%, *p* = 0.001], the medial tibial plateau [−23.8%, CI95: −36.8, −10.9%, *p* < 0.001], and the patella [−29.8%, CI95: −39.5, −20.1%, *p* < 0.001]. Interestingly, the peak proteoglycan content was 21.8% [CI95: 6.2, 37.4%, *p* = 0.009] and 13.2% [CI95: 3.5, 22.8%, *p* = 0.010] greater in the medial tibial plateau and the patella of the CL group rabbits compared to that measured for the CNTRL group rabbits in the middle zone. Finally, the proteoglycan content in the PCM was generally reduced less than in the ECM for the ACLT compared to the CL or CNTRL group rabbits. This trend was observed in both the superficial and middle zone cartilages (Supplementary Table 4).

### 3.3. Relationship of PCM-to-ECM proteoglycan content ratio, pericellular proteoglycan content, and chondrocyte volumetric strain in the superficial zone of cartilage

In the lateral femoral condyle, chondrocytes of the ACLT knees experienced smaller compressive volumetric strain (*p* = 0.05) compared to chondrocytes from the CNTRL group knees. Neither a higher PCM-to-ECM proteoglycan content ratio (*p* = 0.087, Figure 5A) nor a smaller peak proteoglycan content of the PCM (*p* = 0.074, Figure 6A) was observed in the ACLT group to explain the smaller deformation. However, the ACLT group points seemed to form a separate cluster from both the CL and CNTRL groups in the lateral femoral condyle.

**Figure 5.**
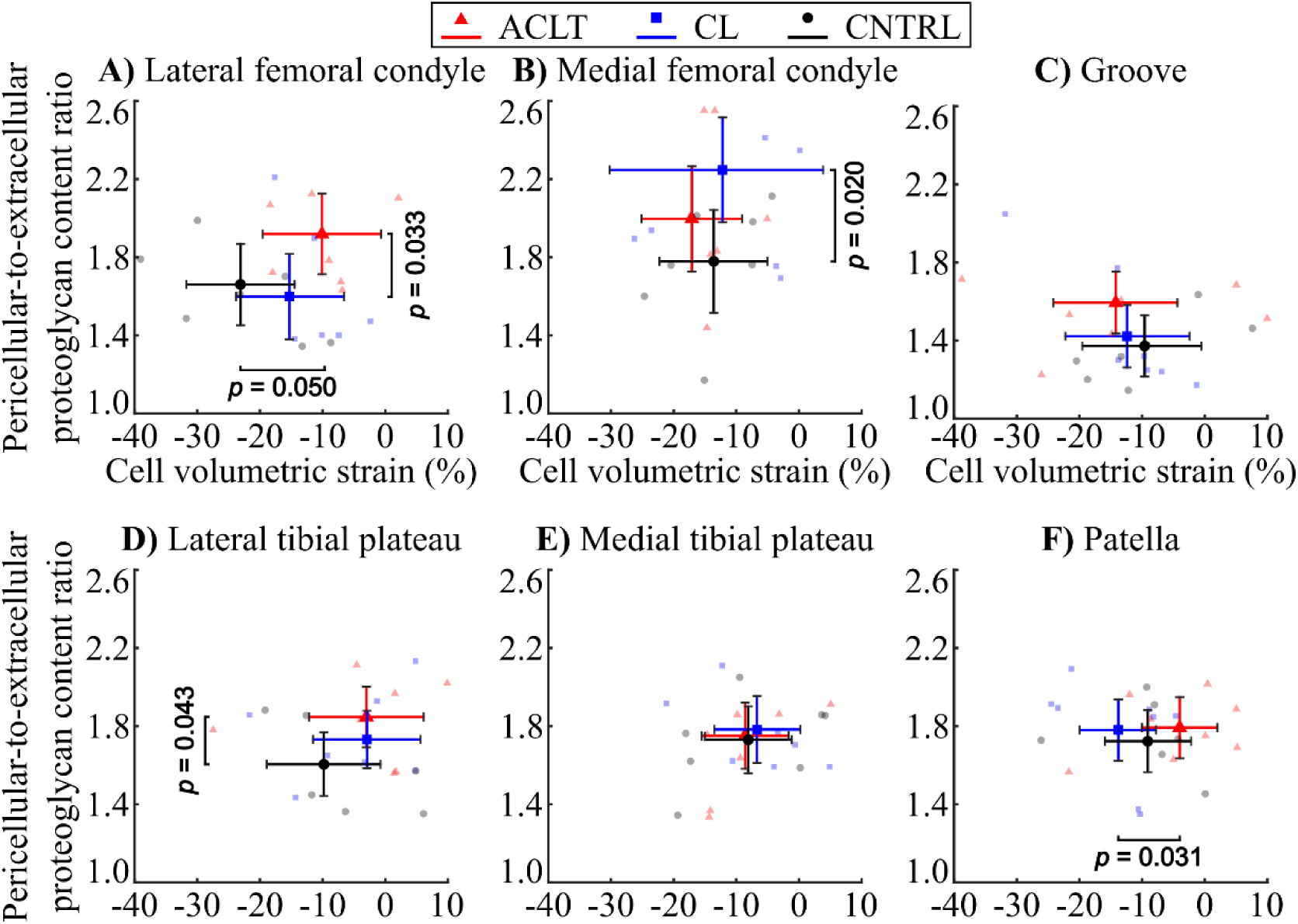
Pericellular-to-extracellular matrix proteoglycan content and cell volumetric strain in the superficial zone cartilage for A) the lateral femoral condyle, B) the medial femoral condyle, C) the lateral femoral groove, D) the lateral tibial plateau, E) the medial tibial plateau, and F) patella. The center markers (triangle, square, circle) represent mean sample-wise values of pericellular proteoglycan content and volumetric cell strain (± 95% CI) for anterior cruciate ligament transection (ACLT, red), contralateral (CL, blue), and the control (CNTRL, black) sample groups.

**Figure 6.**
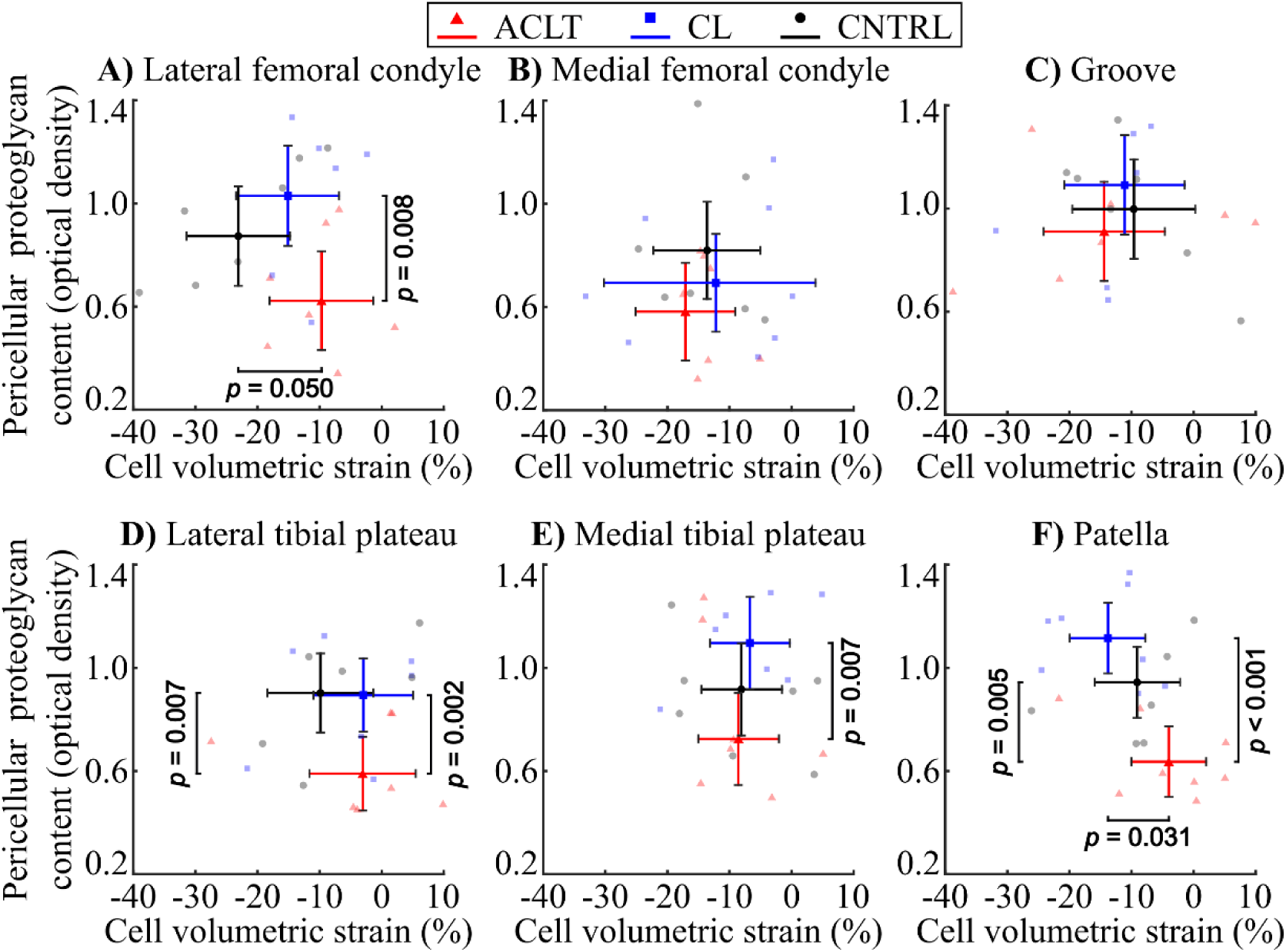
Peak proteoglycan content of the pericellular matrix and cell volumetric strain in the superficial zone cartilage for A) the lateral femoral condyle, B) the medial femoral condyle, C) the lateral femoral groove, D) the lateral tibial plateau, E) the medial tibial plateau, and F) patella. The center markers (triangle, square, circle) represent mean sample-wise values of pericellular proteoglycan content and volumetric cell strain (± 95% CI) for anterior cruciate ligament transection (ACLT, red), contralateral (CL, blue), and the control (CNTRL, black) sample groups.

In the patella, we observed that ACLT knee cartilage had a smaller peak proteoglycan content of the PCM (*p* < 0.001) and smaller compressive volumetric strain (*p* = 0.031) compared to the CL group (Figure 6F). For the same location but compared to the CNTRL group knee cartilage, we also observed that ACLT knee cartilage had a smaller peak proteoglycan content of the PCM (*p* = 0.005) but not a smaller compressive volumetric strain.

## 4. Discussion

We investigated the impact of very early post-traumatic OA, induced by ACLT surgery, on the proteoglycan content in the vicinity of cells in superficial and middle zone rabbit knee cartilages. We observed that the PCM-to-ECM proteoglycan content ratio of the ACLT group was greater compared to the CNTRL group in the middle zone cartilage for the medial femur, femoral groove, and patella. In the superficial zone, the PCM-to-ECM proteoglycan content ratio of the ACLT group was greater compared to the CNTRL group only in the lateral tibial plateau. This suggests that the proteoglycan loss in the PCM was smaller compared to the ECM and that the proteoglycan loss in the PCM might progress more slowly in the middle zone compared to the superficial zone cartilage in early post-traumatic OA.

It has been suggested that the PCM is more resistant to proteoglycan degeneration than the ECM in early OA cartilage [12,34]. The composition of the PCM is distinctly different compared to that of the ECM and, for example, collagen VI and proteoglycans, such as perlecan and biglycan are expressed mainly in the PCM but not the ECM [35–37]. This unique composition of the PCM may explain why the proteoglycan loss was smaller and/or slower in the PCM in the early post-traumatic phase of knee joint OA. For example, heparan sulfate, a key component of perlecan [38], has been reported to inhibit aggrecanase activity [39]. Additionally, biglycan, a small leucine-rich proteoglycan [35], has been shown to have a higher resistance to proteolytic cleavage under inflammatory conditions compared to the main ECM proteoglycan, aggrecan [40].

Overall, the ACLT group had smaller peak proteoglycan content in the PCM compared to the CL and CNTRL group cartilages in the superficial and middle zones. This reduction was more pronounced in the superficial zone cartilage (−32% to −43%) than in the middle zone cartilage (−20% to −30%). We observed similar findings in an earlier study on rabbit cartilage four weeks post-ACLT surgery, where the peak proteoglycan loss in the PCM compared to the CL and CNTRL groups ranged from −28% to −40% in the superficial zone and from −14% to −26% in the middle zone [12]. The observed variation in the proteoglycan content loss of the PCM in different cartilage zones might be due to the different capacity of catabolic activity within the cartilage. It has been suggested that the superficial zone of cartilage has higher levels of ADAMTS-4, MMP-3, and MMP-13 during OA [41,42]. Our gene expression analysis from bulk cartilage [43] indicated an upregulation of MMP-3 and MMP-13 in the ACLT group cartilage. Additionally, IL-6, shown to increase ADAMTS-4 expression [44], was also upregulated. This increased catabolic activity in the superficial compared to the middle zone cartilage might explain the difference in pericellular proteoglycan loss in these two regions.

The CL group had greater peak proteoglycan content in the PCM than the CNTRL group in the middle zone cartilage of the medial tibial plateau and patella. The finding in the medial tibial plateau might be explained by spontaneous degradation (moderately high OARSI score) observed in some CNTRL group rabbits [10]. Generally, differences in the peak proteoglycan content in the superficial and middle zone cartilage were more frequently observed between the ACLT and CL groups than between the ACLT and CNTRL groups. This result could be due to a natural adaptation of proteoglycan content caused by an unloading of the ACLT knees and increased loading of the CL knees [45].

The influence of pericellular proteoglycan content on cell deformation under mechanical loading in the native cartilage was tested for the superficial zone cartilage. In the patellar cartilage, the peak proteoglycan content and the cell volume strain (i.e., volume loss) were smaller in the ACLT group than in the CL group. This result agrees with computational studies, which have shown that with greater PCM proteoglycan content (and greater stiffness), the two orders of magnitude softer cell [33,46] compresses more under cartilage loading, leading to greater cell volume loss [19–22]. However, such results were not observed for the other cartilage surfaces. Since axial cartilage strain during loading did not differ between groups [29], and differences in ECM proteoglycan content did not account for the smaller volumetric cell strain in the ACLT group [29], factors other than decreased proteoglycan content must have contributed to the smaller volumetric cell strain in the ACLT group. One likely factor could be the disorganization or enzymatic cleavage of collagen fibrils near the superficial zone chondrocytes. The organization of collagen fibrils is known to influence the mechanical behavior of both the tissue and chondrocytes, as shown by experimental and computational studies [19,22,47–49], and disorganization of collagen fibrils has been suggested to occur at an early stage of OA, before the loss of collagen in articular cartilage [50,51]. Alternatively, collagen content loss in the PCM could lead to altered force transduction from the cartilage matrix to cells and explain the smaller cell volume strain.

This study has some limitations. The sample groups (8 operated, 8 control rabbits) are small, thus limiting statistical power. Additionally, the proteoglycan content was not analyzed from the same cells as the cell deformation [29]. To account for this difference, cells (*n* = 11 per zone from each location and sample) were analyzed via digital densitometry from the same region where the deformation analysis was done. Also, the proteoglycan content was indirectly estimated from Safranin-O-stained sections. Safranin-O binds stoichiometrically to the negative charges of the glycosaminoglycan chains of proteoglycans [31]. Thus, the measured optical density reflects fixed charge density in cartilage and provides a surrogate measure for spatial proteoglycan content [32]. Finally, the surgical transection of the ACL in the rabbit model does not perfectly mimic the uncontrolled ACL rupture in humans caused by (often sporting) accidents. However, changes in the physical properties of cartilage between surgical ACLT and injury-induced tearing of the ACL have been shown to be similar [26]. Naturally, the small but distinct differences between human and rabbit knee joint shapes, cartilage structure, and cartilage biomechanics must be kept in mind when translating our results to those associated with human ACL tears caused by accidents.

Based on the results of this study, we conclude that two weeks post-ACLT the proteoglycan content loss is smaller in the PCM compared to the ECM, and this is more prevalent in the middle zone compared to the superficial zone of cartilage. Moreover, these changes in the superficial zone did not have a clear association to the observed smaller chondrocyte volumetric strains during controlled compressive loading of the cartilage tissue.

## Acknowledgements

This work was supported by the State Research Funding for university-level health research (Kuopio University Hospital) Wellbeing Services County of North Savo (#5654244, #5654265), Research Council of Finland (#363459, #354916), Sigrid Jusélius Foundation, Novo Nordisk Foundation (#NNF21OC0065373), Maire Lisko Foundation, Strategic Funding of the University of Eastern Finland, Kuopio University Foundation, Saastamoinen Foundation, Päivikki ja Sakari Sohlberg Foundation (#240074), Finnish Cultural Foundation North Savo Regional Fund (#65171624), The Arthritis Society of Canada, The Canadian Institutes of Health Research, and the Nigg Chair for mobility and longevity. The authors would like to thank Santtu Mikkonen, PhD, University of Eastern Finland, for consultation in the statistical analysis, Andrew Sawatsky, MSc, University of Calgary, for ACL surgeries and sample preparation, and Tarja Huhta, University of Oulu, for the sample preparations.

## References

[1] N. Wang, Review of cellular mechanotransduction, J Phys D Appl Phys 50 (2017) 233002. 10.1088/1361-6463/aa6e18.

[2] A.J. Sophia Fox, A. Bedi, S.A. Rodeo, The Basic Science of Articular Cartilage: Structure, Composition, and Function, Sports Health: A Multidisciplinary Approach 1 (2009) 461–468. 10.1177/1941738109350438.

[3] H. Yoshida, T. Kojima, K. Kurokouchi, S. Takahashi, H. Hanamura, M. Kojima, A.R. Poole, N. Ishiguro, Relationship between pre-radiographic cartilage damage following anterior cruciate ligament injury and biomarkers of cartilage turnover in clinical practice: a cross-sectional observational study, Osteoarthritis Cartilage 21 (2013) 831– 838. 10.1016/j.joca.2013.03.009.

[4] H. Long, Q. Liu, H. Yin, K. Wang, N. Diao, Y. Zhang, J. Lin, A. Guo, Prevalence Trends of Site-Specific Osteoarthritis From 1990 to 2019: Findings From the Global Burden of Disease Study 2019, Arthritis & Rheumatology 74 (2022) 1172–1183. 10.1002/art.42089.

[5] L. Punzi, P. Galozzi, R. Luisetto, M. Favero, R. Ramonda, F. Oliviero, A. Scanu, Post-traumatic arthritis: overview on pathogenic mechanisms and role of inflammation, RMD Open 2 (2016) e000279. 10.1136/rmdopen-2016-000279.

[6] Y. He, Z. Li, P.G. Alexander, B.D. Ocasio-Nieves, L. Yocum, H. Lin, R.S. Tuan, Pathogenesis of Osteoarthritis: Risk Factors, Regulatory Pathways in Chondrocytes, and Experimental Models, Biology (Basel) 9 (2020) 194. 10.3390/biology9080194.

[7] Y. Sun, P. Leng, P. Guo, H. Gao, Y. Liu, C. Li, Z. Li, H. Zhang, G protein coupled estrogen receptor attenuates mechanical stress-mediated apoptosis of chondrocyte in osteoarthritis via suppression of Piezo1, Molecular Medicine 27 (2021) 96. 10.1186/s10020-021-00360-w.

[8] L. Ramage, G. Nuki, D.M. Salter, Signalling cascades in mechanotransduction: cell– matrix interactions and mechanical loading, Scand J Med Sci Sports 19 (2009) 457–469. 10.1111/j.1600-0838.2009.00912.x.

[9] G. Venn, M.E.J. Billingham, T.E. Hardingham, Increased proteoglycan synthesis in cartilage in experimental canine osteoarthritis does not reflect a permanent change in chondrocyte phenotype, Arthritis Rheum 38 (1995) 525–532. 10.1002/art.1780380410.

[10] L. Huang, I. Riihioja, P. Tanska, S. Ojanen, S. Palosaari, H. Kröger, S.J. Saarakkala, W. Herzog, R.K. Korhonen, M.A.J. Finnilä, Early changes in osteochondral tissues in a rabbit model of post-traumatic osteoarthritis, Journal of Orthopaedic Research 39 (2021) 2556–2567. 10.1002/jor.25009.

[11] T.M. Quinn, P. Schmid, E.B. Hunziker, A.J. Grodzinsky, Proteoglycan deposition around chondrocytes in agarose culture: construction of a physical and biological interface for mechanotransduction in cartilage., Biorheology 39 (2002) 27–37. http://www.ncbi.nlm.nih.gov/pubmed/12082264.

[12] S.P. Ojanen, M.A.J. Finnilä, A.E. Reunamo, A.P. Ronkainen, S. Mikkonen, W. Herzog, S. Saarakkala, R.K. Korhonen, Site-specific glycosaminoglycan content is better maintained in the pericellular matrix than the extracellular matrix in early post-traumatic osteoarthritis, PLoS One 13 (2018) e0196203. 10.1371/journal.pone.0196203.

[13] H. Akkiraju, A. Nohe, Role of Chondrocytes in Cartilage Formation, Progression of Osteoarthritis and Cartilage Regeneration, J Dev Biol 3 (2015) 177–192. 10.3390/jdb3040177.

[14] F. Guilak, R.J. Nims, A. Dicks, C.-L. Wu, I. Meulenbelt, Osteoarthritis as a disease of the cartilage pericellular matrix, Matrix Biology 71–72 (2018) 40–50. 10.1016/j.matbio.2018.05.008.

[15] E.M. Darling, R.E. Wilusz, M.P. Bolognesi, S. Zauscher, F. Guilak, Spatial Mapping of the Biomechanical Properties of the Pericellular Matrix of Articular Cartilage Measured In Situ via Atomic Force Microscopy, Biophys J 98 (2010) 2848–2856. 10.1016/j.bpj.2010.03.037.

[16] T.M. Quinn, A.A. Maung, A.J. Grodzinsky, E.B. Hunziker, J.D. Sandy, Physical and Biological Regulation of Proteoglycan Turnover around Chondrocytes in Cartilage Explants: Implications for Tissue Degradation and Repair, Ann N Y Acad Sci 878 (1999) 420–441. 10.1111/j.1749-6632.1999.tb07700.x.

[17] F. Guilak, L.G. Alexopoulos, M.L. Upton, I. Youn, J.B. Choi, L. Cao, L.A. Setton, M.A. Haider, The Pericellular Matrix as a Transducer of Biomechanical and Biochemical Signals in Articular Cartilage, Ann N Y Acad Sci 1068 (2006) 498–512. 10.1196/annals.1346.011.

[18] J.B. Choi, I. Youn, L. Cao, H.A. Leddy, C.L. Gilchrist, L.A. Setton, F. Guilak, Zonal changes in the three-dimensional morphology of the chondron under compression: The relationship among cellular, pericellular, and extracellular deformation in articular cartilage, J Biomech 40 (2007) 2596–2603. 10.1016/j.jbiomech.2007.01.009.

[19] P. Tanska, S.M. Turunen, S.K. Han, P. Julkunen, W. Herzog, R.K. Korhonen, Superficial Collagen Fibril Modulus and Pericellular Fixed Charge Density Modulate Chondrocyte Volumetric Behaviour in Early Osteoarthritis, Comput Math Methods Med 2013 (2013) 1–14. 10.1155/2013/164146.

[20] P. Tanska, M.S. Venäläinen, A. Erdemir, R.K. Korhonen, A multiscale framework for evaluating three-dimensional cell mechanics in fibril-reinforced poroelastic tissues with anatomical cell distribution – Analysis of chondrocyte deformation behavior in mechanically loaded articular cartilage, J Biomech 101 (2020) 109648. 10.1016/j.jbiomech.2020.109648.

[21] R.K. Korhonen, W. Herzog, Depth-dependent analysis of the role of collagen fibrils, fixed charges and fluid in the pericellular matrix of articular cartilage on chondrocyte mechanics, J Biomech 41 (2008) 480–485. 10.1016/j.jbiomech.2007.09.002.

[22] A.P. Ronkainen, P. Tanska, J.M. Fick, W. Herzog, R.K. Korhonen, Interrelationship of cartilage composition and chondrocyte mechanics after a partial meniscectomy in the rabbit knee joint – Experimental and numerical analysis, J Biomech 83 (2019) 65–75. 10.1016/j.jbiomech.2018.11.024.

[23] D.R. Chery, B. Han, Q. Li, Y. Zhou, S.-J. Heo, B. Kwok, P. Chandrasekaran, C. Wang, L. Qin, X.L. Lu, D. Kong, M. Enomoto-Iwamoto, R.L. Mauck, L. Han, Early changes in cartilage pericellular matrix micromechanobiology portend the onset of post-traumatic osteoarthritis, Acta Biomater 111 (2020) 267–278. 10.1016/j.actbio.2020.05.005.

[24] A. Linus, M. Ebrahimi, M.J. Turunen, S. Saarakkala, A. Joukainen, H. Kröger, A. Koistinen, M.A.J. Finnilä, I.O. Afara, M.E. Mononen, P. Tanska, R.K. Korhonen, High-resolution infrared microspectroscopic characterization of cartilage cell microenvironment, Acta Biomater 134 (2021) 252–260. 10.1016/j.actbio.2021.08.001.

[25] R.D. Altman, D.D. Dean, Osteoarthritis research: Animal models, Semin Arthritis Rheum 19 (1990) 21–25. 10.1016/0049-0172(90)90081-P.

[26] R.L. Sah, A.S. Yang, A.C. Chen, J.J. Hant, R.B. Halili, M. Yoshioka, D. Amiel, R.D. Coutts, Physical properties of rabbit articular cartilage after transection of the anterior cruciate ligament, Journal of Orthopaedic Research 15 (1997) 197–203. 10.1002/jor.1100150207.

[27] S.M. Turunen, S.-K. Han, W. Herzog, R.K. Korhonen, Cell deformation behavior in mechanically loaded rabbit articular cartilage 4 weeks after anterior cruciate ligament transection, Osteoarthritis Cartilage 21 (2013) 505–513. 10.1016/j.joca.2012.12.001.

[28] J.T.A. Mäkelä, Z.S. Rezaeian, S. Mikkonen, R. Madden, S.-K. Han, J.S. Jurvelin, W. Herzog, R.K. Korhonen, Site-dependent changes in structure and function of lapine articular cartilage 4 weeks after anterior cruciate ligament transection, Osteoarthritis Cartilage 22 (2014) 869–878. 10.1016/j.joca.2014.04.010.

[29] S.P. Ojanen, M.A.J. Finnilä, J.T.A. Mäkelä, K. Saarela, E. Happonen, W. Herzog, S. Saarakkala, R.K. Korhonen, Anterior cruciate ligament transection of rabbits alters composition, structure and biomechanics of articular cartilage and chondrocyte deformation 2 weeks post-surgery in a site-specific manner, J Biomech 98 (2020) 109450. 10.1016/j.jbiomech.2019.109450.

[30] S.-K. Han, P. Colarusso, W. Herzog, Confocal microscopy indentation system for studying in situ chondrocyte mechanics, Med Eng Phys 31 (2009) 1038–1042. 10.1016/j.medengphy.2009.05.013.

[31] I. Kiviranta, J. Jurvelin, A.-M. Säämänen, H.J. Helminen, Microspectrophotometric quantitation of glycosaminoglycans in articular cartilage sections stained with Safranin O, Histochemistry 82 (1985) 249–255. 10.1007/BF00501401.

[32] K. Király, T. Lapveteläinen, J. Arokoski, K. Törrönen, L. Módis, I. Kiviranta, H.J. Helminen, Application of selected cationic dyes for the semiquantitative estimation of glycosaminoglycans in histological sections of articular cartilage by microspectrophotometry, Histochem J 28 (1996) 577–590. 10.1007/BF02331378.

[33] R.E. Wilusz, J. Sanchez-Adams, F. Guilak, The structure and function of the pericellular matrix of articular cartilage, Matrix Biology 39 (2014) 25–32. 10.1016/j.matbio.2014.08.009.

[34] R.E. Wilusz, F. Guilak, High resistance of the mechanical properties of the chondrocyte pericellular matrix to proteoglycan digestion by chondroitinase, aggrecanase, or hyaluronidase, J Mech Behav Biomed Mater 38 (2014) 183–197. 10.1016/j.jmbbm.2013.09.021.

[35] E. Kavanagh, D.E. Ashhurst, Development and Aging of the Articular Cartilage of the Rabbit Knee Joint: Distribution of Biglycan, Decorin, and Matrilin-1, Journal of Histochemistry & Cytochemistry 47 (1999) 1603–1615. 10.1177/002215549904701212.

[36] R. Gomes, C. Kirn-Safran, M.C. Farach-Carson, D.D. Carson, Perlecan: an important component of the cartilage pericellular matrix., J Musculoskelet Neuronal Interact 2 (2002) 511–6.

[37] C.A. Poole, S. Ayad, R.T. Gilbert, Chondrons from articular cartilage: V.* Immunohistochemical evaluation of type VI collagen organisation in isolated chondrons by light, confocal and electron microscopy, J Cell Sci 103 (1992) 1101–1110. 10.1242/jcs.103.4.1101.

[38] M.A. Gubbiotti, T. Neill, R. V. Iozzo, A current view of perlecan in physiology and pathology: A mosaic of functions, Matrix Biology 57–58 (2017) 285–298. 10.1016/j.matbio.2016.09.003.

[39] S.E. Munteanu, M.Z. Ilic, C.J. Handley, Highly sulfated glycosaminoglycans inhibit aggrecanase degradation of aggrecan by bovine articular cartilage explant cultures, Matrix Biology 21 (2002) 429–440. 10.1016/S0945-053X(02)00034-3.

[40] R. Sztrolovics, R.J. White, A.R. Poole, J.S. Mort, P.J. Roughley, Resistance of small leucine-rich repeat proteoglycans to proteolytic degradation during interleukin-1-stimulated cartilage catabolism., Biochem J 339 ( Pt 3) (1999) 571–7.

[41] N. Fukui, Y. Miyamoto, M. Nakajima, Y. Ikeda, A. Hikita, H. Furukawa, H. Mitomi, N. Tanaka, Y. Katsuragawa, S. Yamamoto, M. Sawabe, T. Juji, T. Mori, R. Suzuki, S. Ikegawa, Zonal gene expression of chondrocytes in osteoarthritic cartilage, Arthritis Rheum 58 (2008) 3843–3853. 10.1002/art.24036.

[42] K.S.C. Cheung, K. Hashimoto, N. Yamada, H.I. Roach, Expression of ADAMTS-4 by chondrocytes in the surface zone of human osteoarthritic cartilage is regulated by epigenetic DNA de-methylation, Rheumatol Int 29 (2009) 525–534. 10.1007/s00296-008-0744-z.

[43] M.A. Finnilä, S. Ojanen, S. Saarakkala, C. Hewitt, W. Herzog, P. Nieminen, D.A. Hart, R.K. Korhonen, Increased cartilage remodelling and impaired chondrocyte mechanotransduction in early post-traumatic osteoartritis, Osteoarthritis Cartilage 25 (2017) S67–S68. 10.1016/j.joca.2017.02.122.

[44] N. Sahu, H.J. Viljoen, A. Subramanian, Continuous low-intensity ultrasound attenuates IL-6 and TNFα-induced catabolic effects and repairs chondral fissures in bovine osteochondral explants, BMC Musculoskelet Disord 20 (2019) 193. 10.1186/s12891-019-2566-4.

[45] M. Tammi, A.-M. Säämänen, A. Jauhiainen, O. Malminen, I. Kiviranta, H. Helminen, Proteoglycan Alterations in Rabbit Knee Articular Cartilage Following Physical Exercise and Immobilization, Connect Tissue Res 11 (1983) 45–55. 10.3109/03008208309015010.

[46] C. Florea, P. Tanska, M.E. Mononen, C. Qu, M.J. Lammi, M.S. Laasanen, R.K. Korhonen, A combined experimental atomic force microscopy-based nanoindentation and computational modeling approach to unravel the key contributors to the time-dependent mechanical behavior of single cells, Biomech Model Mechanobiol 16 (2017) 297–311. 10.1007/s10237-016-0817-y.

[47] S.M. Turunen, M.J. Lammi, S. Saarakkala, S.-K. Han, W. Herzog, P. Tanska, R.K. Korhonen, The effect of collagen degradation on chondrocyte volume and morphology in bovine articular cartilage following a hypotonic challenge, Biomech Model Mechanobiol 12 (2013) 417–429. 10.1007/s10237-012-0409-4.

[48] R.K. Korhonen, M. Wong, J. Arokoski, R. Lindgren, H.J. Helminen, E.B. Hunziker, J.S. Jurvelin, Importance of the superficial tissue layer for the indentation stiffness of articular cartilage, Med Eng Phys 24 (2002) 99–108. 10.1016/S1350-4533(01)00123-0.

[49] M.R.J. Huttu, J. Puhakka, J.T.A. Mäkelä, Y. Takakubo, V. Tiitu, S. Saarakkala, Y.T. Konttinen, I. Kiviranta, R.K. Korhonen, Cell–tissue interactions in osteoarthritic human hip joint articular cartilage, Connect Tissue Res 55 (2014) 282–291. 10.3109/03008207.2014.912645.

[50] E.K. Moo, M. Ebrahimi, S.C. Sibole, P. Tanska, R.K. Korhonen, The intrinsic quality of proteoglycans, but not collagen fibres, degrades in osteoarthritic cartilage, Acta Biomater 153 (2022) 178–189. 10.1016/j.actbio.2022.09.002.

[51] S. Saarakkala, P. Julkunen, P. Kiviranta, J. Mäkitalo, J.S. Jurvelin, R.K. Korhonen, Depth-wise progression of osteoarthritis in human articular cartilage: investigation of composition, structure and biomechanics, Osteoarthritis Cartilage 18 (2010) 73–81. 10.1016/j.joca.2009.08.003.

