## Supplementary for "Pericellular Matrix Proteoglycan Content is Less Affected in The Mid-Zone Cartilage in Early Post-Traumatic Osteoarthritis"

Supplementary Material

**Section A. Supplementary methods**

**A.1. Animal model**

A unilateral anterior cruciate ligament transection (ACLT) was performed in eight skeletally mature female New Zealand white rabbits (Oryctolagus cuniculus, age 12 months at the time of surgery, weight 4.44 ± 0.45 kg [mean ± standard deviation]) where the operated knee joint was chosen randomly for each rabbit to avoid bias (Ojanen et al., 2020). Rabbits were euthanized two weeks post-surgery under anesthesia, and both operated (ACLT) and contralateral (CL) knee joints were harvested. Randomly selected knee joints (left or right leg) from eight age-matched and unoperated healthy female rabbits were collected for the control group (CNTRL, weight 4.57 ± 0.35 kg). Rabbits of the same sex were used to negate gender-related differences in osteoarthritis (OA) development and progression. All procedures were approved by the Committee on Animal Ethics at the University of Calgary and were carried out according to their guidelines (research permit #AC11-0035).

**A.2. In-situ two-photon confocal microscopy and indentation testing** (Ojanen et al., 2020)

The collected osteochondral samples were immersed in Dulbecco Modified Eagle Medium (DMEM, high glucose, HEPES, no phenol red, 21063-029, Grand Island, NY, USA) with Dextran (excitation 488 nm, emission 515 nm, D34682, Thermo Scientific, Eugene, Oregon, USA) and Propidium iodide (P3566, Thermo Scientific, Eugene, Oregon, USA) stains for 4 h at 4ºC. The extracellular matrix (ECM) was stained with Dextran to identify chondrocytes and Propidium iodide was used to label the dead cells. Excess stains were removed by rinsing the samples twice with DMEM for 2 minutes. After rinsing, the samples were fixed in place in a custom-made sample holder with dental cement. The cement was left to cure for 15-20 minutes. For the whole duration of the experiment, the samples were hydrated with DMEM.

After sample fixation, confocal microscopy imaging with *in-situ* indentation testing was conducted according to a previously validated protocol (Han et al., 2009). The testing setup allows confocal imaging with simultaneous indentation loading of the sample. In the measurement setup, a cylindrical glass indenter (diameter 2 mm) is placed under a dual-photon excitation microscope (Chameleon XR infrared laser, Coherent Inc., USA). A 40x magnification water immersion objective (NA: 0.8, Zeiss Inc., Germany) is used for a spatial resolution of 0.41×0.41×0.50 µm^3^ (x, y, and z directions). Measurements were performed at room temperature (21 degrees Celsius) and the load was applied by compressing the sample against the glass indenter. To achieve proper contact between cartilage and the indenter, a preload of 0.1—0.2 MPa was applied after which the tissue was allowed to relax for 20 minutes. After relaxation, an image stack was collected for pre-load cell morphology. Following the image collection, indentation with a 10 µm/s ramp rate was applied to the cartilage until a terminal load of 2 MPa was reached. Afterward, the tissue was again allowed to relax for 20 minutes while the indenter displacement was kept constant. A second image stack was then collected for analysis of the after-load cell morphology. Indentation was performed at the most prominent site for the femoral groove, femoral condyles, and patella and in the geometric sagittal plane center for the tibial plateaus (Mäkelä et al., 2014).

The cell morphology of roughly 40-70 viable superficial chondrocytes from each cartilage location of each measurement group was measured, both before and after loading. The volume, surface area, height, width, and depth of the chondrocytes were measured and the relative change in these parameters before and after loading was analyzed. Width and height were defined as the major and minor axes of the cell in the x-y-plane, respectively, and depth as the thickness of the cell in the z-direction. Deformation due to loading was analyzed for all cell dimensions, cell volume, and surface area. Additionally, the axial and transversal engineering strains of superficial zone cartilage ECM were analyzed via identified cell pairs (4 cell pairs per sample).

**A.3. Sample preparation**

After the two-photon microscopy, the samples were fixed in formalin, decalcified using ethylenediaminetetraacetic (EDTA) acid, dehydrated in ascending alcohol series, and embedded in paraffin. Three histological sections (3 μm thick) per sample were prepared and stained with Safranin-O from each joint location and each sample group. In addition, three unstained sections (5 µm thick) per sample were prepared for polarized light microscopy at the same locations. The superficial and middle zones of each cartilage location were defined based on the average depth-wise collagen orientation angle profile of the CNTRL group using polarized light microscopy (Ojanen et al., 2020).

**A.4. Digital densitometry**

Digital densitometry performed on the Safranin-O-stained histological sections was used to spatially evaluate the amount of proteoglycans (Kiviranta et al., 1985; Ojanen et al., 2018) of the chondrocyte microenvironment, which consists of the pericellular matrix (PCM) and surrounding ECM. In digital densitometry, the optical density is measured due to Safranin-O binding to the negative charges of cartilage, providing a surrogate measure of the proteoglycan content and fixed charge density in the cartilage tissue samples (Király et al., 1996; Kiviranta et al., 1985; Ojanen et al., 2018). The digital densitometry was performed with a light microscope (Microphot FXA; Nikon, Tokyo, Japan) connected to a CCD digital camera (ORCA-ER; Hamamatsu Photonics K.K, Hamamatsu City, Japan). Monochromatic light and 40x magnification (NA: 0.7, λ: 492 ± 5 nm) were used for a spatial resolution of 0.43×0.43 μm^2^. Images were calibrated into optical density images with neutral density filters (optical density values of air, 0.0, 0.3, 0.6, 1.0, 1.3, 1.6, 2.0, 2.3, 2.6, and 3.0) (Schott, Mainz, Germany).

Based on light microscopy images of the Safranin-O-stained sections, cells were selected from the regions of the cartilage surfaces where the cell morphology was analyzed. However, the cells were not necessarily the same ones measured with two-photon confocal microscopy. On average, 72 cells were selected to be imaged per zone in each location and sample group. The total number of cells is shown in Supplementary Table 1. Safranin-O binds stoichiometrically to the negatively charged glycosaminoglycans in tissue. Thus, measurement of the optical density from Safranin-O-stained sections allows for the estimation of proteoglycan content (Kiviranta et al., 1985).

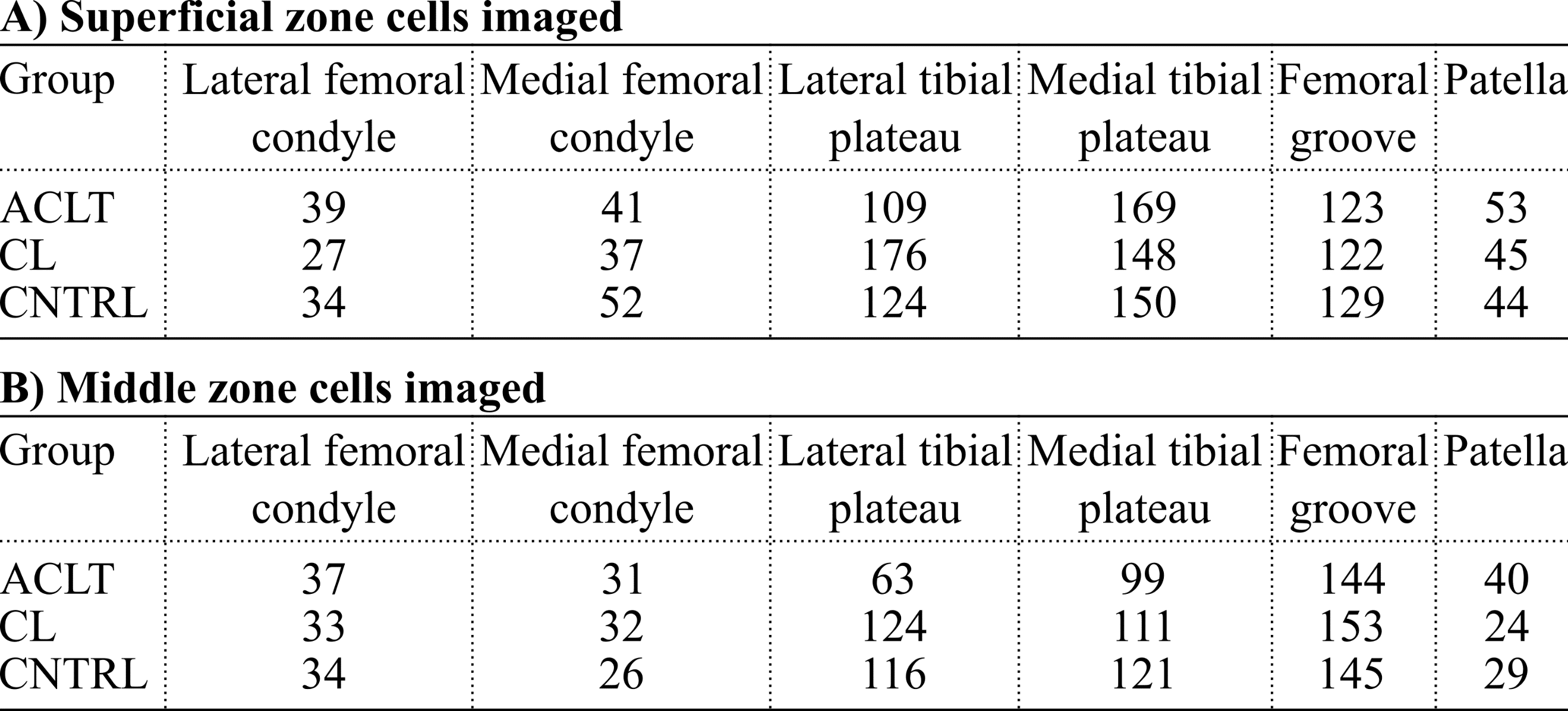

***Supplementary Table 1.*** *The number of cells imaged with digital densitometry in A) the superficial zone cartilage and B) the middle zone cartilage. The amounts are shown separately for the anterior cruciate ligament transected (ACLT), contralateral (CL) and control (CNTRL) group knees (rows) for each analyzed joint surface (columns).*

**A.5. Analysis of the cell microenvironment**

The cells were manually segmented using light microscopy images (Supplementary Figure 1). The absolute proteoglycan content profiles of the chondrocyte microenvironment were averaged from a rectangular region (height: 6 μm) that laterally extended 20 μm from the cell border towards the ECM (Figure 1F). The rectangular region was set parallel to the major axis of the imaged chondrocytes. This was done because superficial zone cells are, on average, oriented parallel to the cartilage surface. Thus, for each measured cell we had absolute proteoglycan content profiles from both sides of the cell, labeled either as “left” or “right” profiles that were separated by the segmented cell.

***Supplementary Figure 1.*** *Cell segmentation from light microscopy images. A) The cell border was determined as the inner interface between stained and non-stained regions of the cell, where the strongly stained rim region surrounding the cell was presumed to be the pericellular matrix (Ojanen et al., 2018; Ronkainen et al., 2016). B) The cell was segmented based on the determined interface and removed from the spatial proteoglycan content (optical density) maps obtained via digital densitometry.*

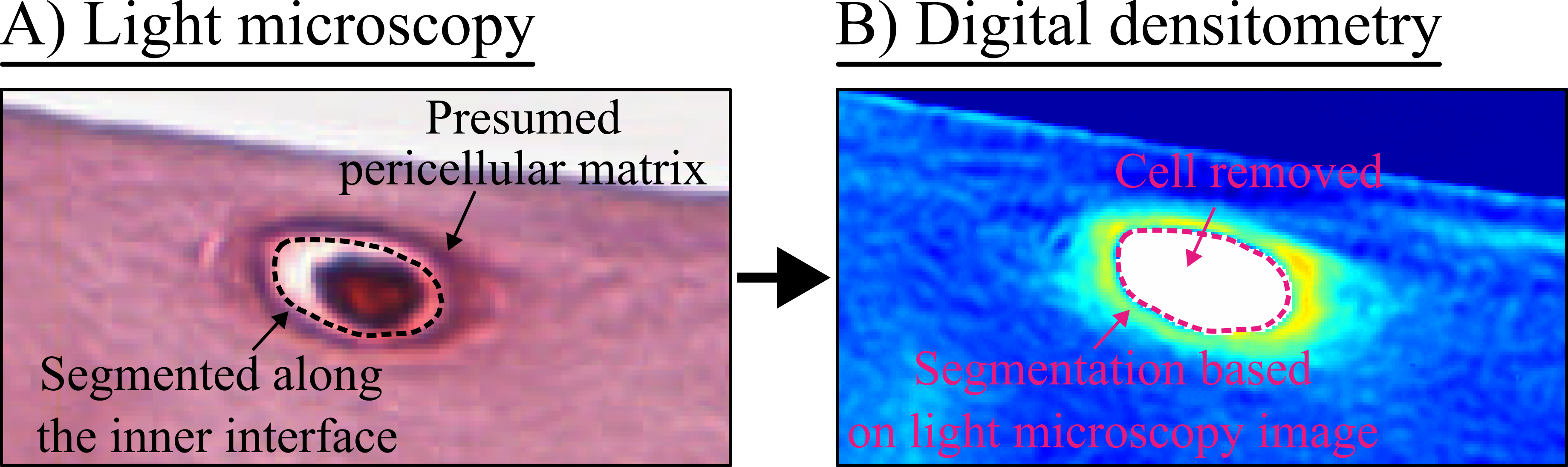

From the absolute profiles, we first analyzed the peak value within 5 µm of the cell edge. This 5 µm thick region presumably contains the PCM and the peak value was used to represent proteoglycan content of the PCM. Second, we calculated normalized proteoglycan content profiles by dividing each absolute profile by the point 20 μm from the cell edge. The normalization highlights changes in the proteoglycan content of the PCM and territorial ECM relative to the interterritorial ECM. Additionally, the value 20 µm from the cell edge was used to represent the proteoglycan content of the ECM. Lastly, the peak value from the normalized profiles (labeled as the PCM/ECM ratio) was calculated. The PCM/ECM ratio was analyzed separately from the normalized proteoglycan content profiles to account for variation in the peak location between individual cells (due to possible differences in the PCM thickness between cells). For example, we observed that the peak value was closer to the cell edge in the ACLT group knees compared to the CL and CNTRL group knees (Supplementary Tables 2 and 3).

***Supplementary Table 2.*** *Cell distance from the cartilage surface and distance to the peak proteoglycan content value from the cell edge, and their 95% confidence intervals (95% CI) for superficial and middle zone chondrocytes in the anterior cruciate ligament transection (ACLT), contralateral (CL), and separate healthy control (CNTRL) sample groups. Cell distance from the cartilage surface was calculated from optical images for each cell as the distance of the cell to the surface relative to the distance between the cartilage surface and the subchondral bone. Distance to the peak value indicates the distance from the cell edge to the peak proteoglycan content value in the proteoglycan content profile.*

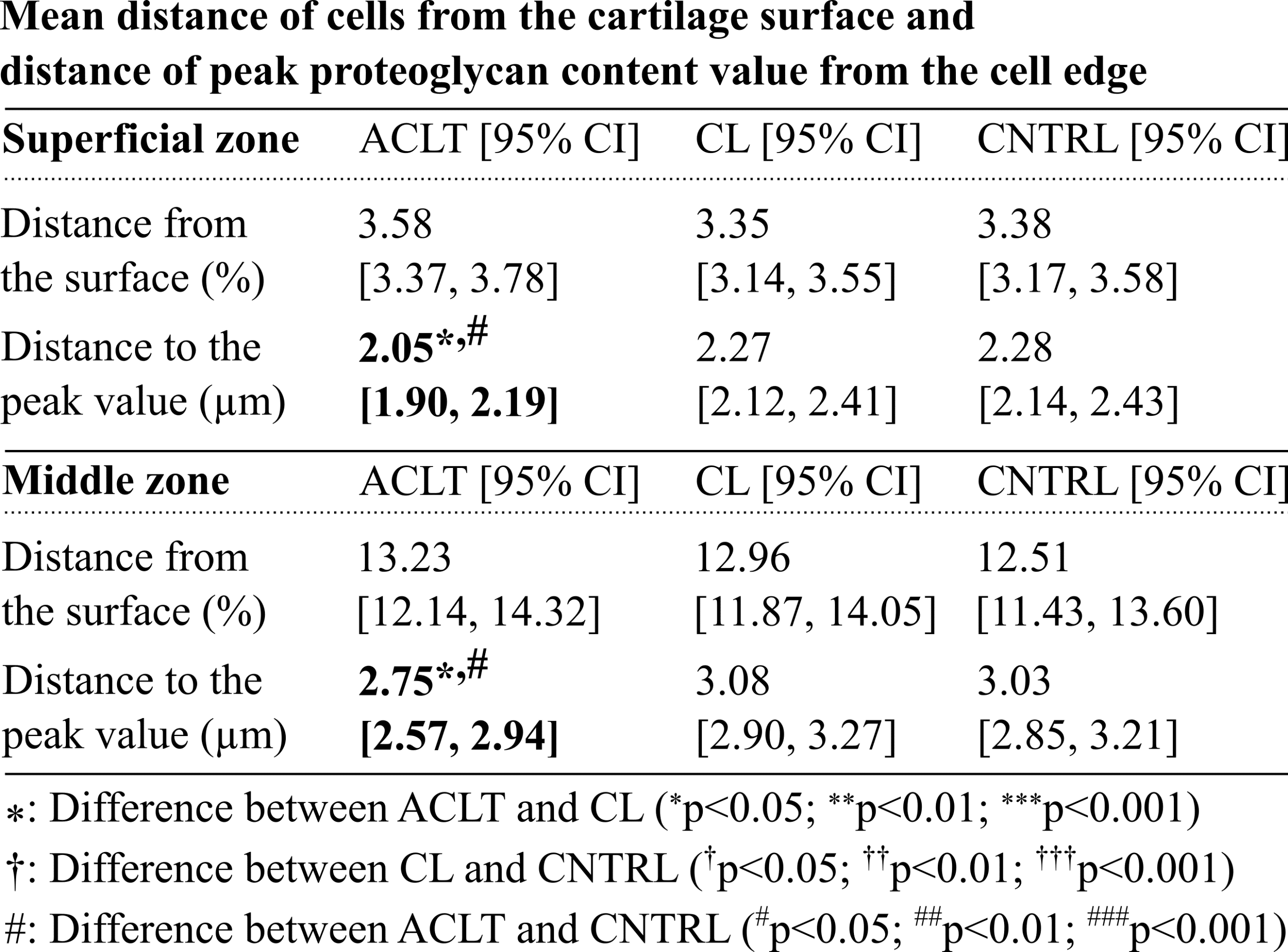

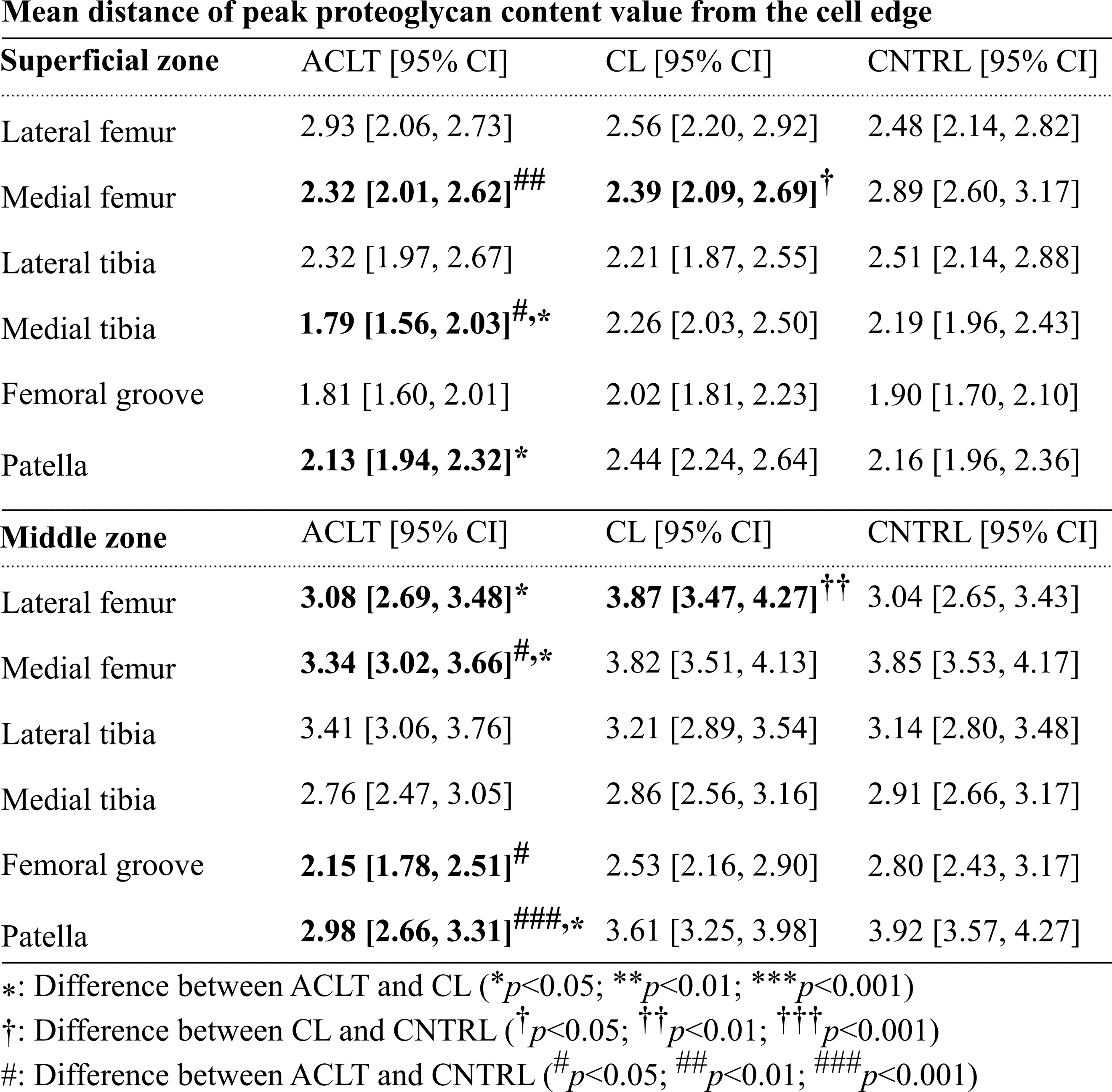

***Supplementary Table 3.*** *Mean distance to the peak proteoglycan content (optical density) value from the cell edge and 95% confidence intervals (CIs) in the superficial and middle zone cartilage for all analyzed locations: lateral and medial femoral condyle, lateral and medial tibial plateau, femoral groove, and patella. ACLT, anterior cruciate ligament transection; CL, contralateral; CNTRL, the healthy control group.*

**Section B. Supplementary results**

**B.1. Absolute proteoglycan content of the cell microenvironment**

In the superficial zone of cartilage, we observed differences in the absolute proteoglycan content profiles where a difference in peak proteoglycan content was also observed (Figure 2A, Supplementary Figure 2A). Absolute proteoglycan content was lower in the ACLT group knees compared to the CL group knees for each joint surface except the medial femoral condyle. Additionally, in the femoral groove, the difference was observed only in the ECM. While in the lateral femoral condyle, tibial plateaus, and patella proteoglycan content of the ACLT groups knees were lower near the cell and this extended to the ECM. In the lateral tibial plateau and patella, the absolute proteoglycan content was lower in the ACLT group knees compared to the CNTRL group knees for the whole 20 µm distance that was analyzed.

As before, in the middle zone cartilage, we observed differences in the absolute proteoglycan content profiles between the groups in the pericellular and territorial regions at each location where we had observed a difference in the peak proteoglycan content (Figure 2B, Supplementary Figure 2B). Absolute proteoglycan content was lower in the ACLT group knees compared to the CNTRL group knees for the medial femoral condyle and patella. Compared to the CL group knees, proteoglycan content was lower in the ACLT group knees for each location. However, in the lateral tibial plateau, this difference was observed only near the cell edge. Interestingly, proteoglycan content was higher in the CL group knees compared to the CNTRL group knees for the lateral femoral condyle, medial tibial plateau, and patella.

Overall, the difference in the ACLT group knees compared to the CL and CNTRL group knees was higher in the ECM (20 µm from the cell edge) than in the PCM (Supplementary Table 4). On average, the difference between the groups was 7.5% higher in the ECM compared to the PCM.

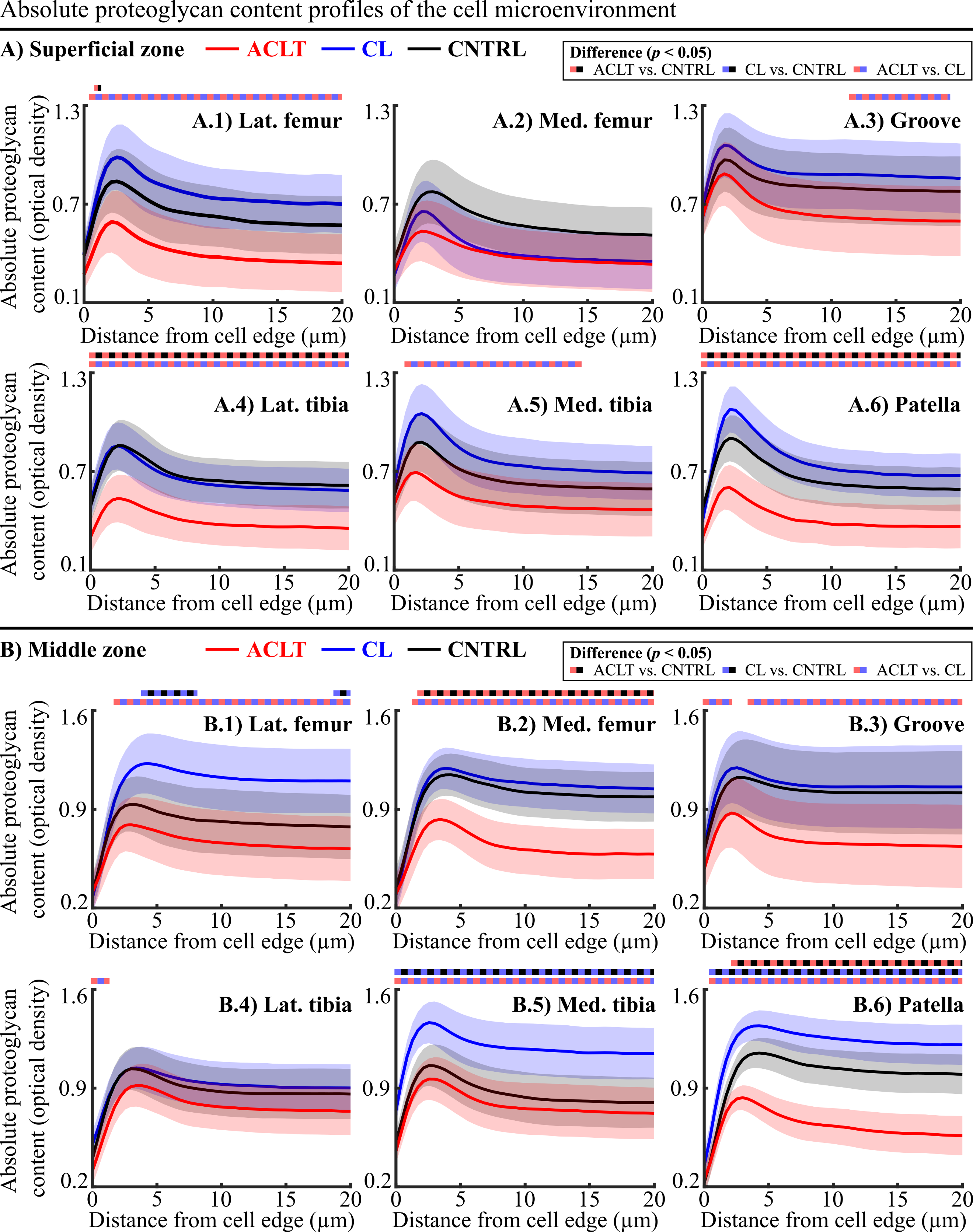

***Supplementary Figure 2.*** *Absolute proteoglycan content profiles for the A) superficial and B) middle zone cell microenvironment in the 1) lateral femoral condyle, 2) medial femoral condyle, 3) lateral femoral groove, 4) lateral tibial plateau, 5) medial tibial plateau, and 6) patella. The mean profiles are drawn with solid lines for anterior cruciate ligament transection (ACLT, red), contralateral (CL, blue), and the separate healthy control (CNTRL, black) sample groups. Shaded areas represent each mean profile's 95% confidence intervals.*

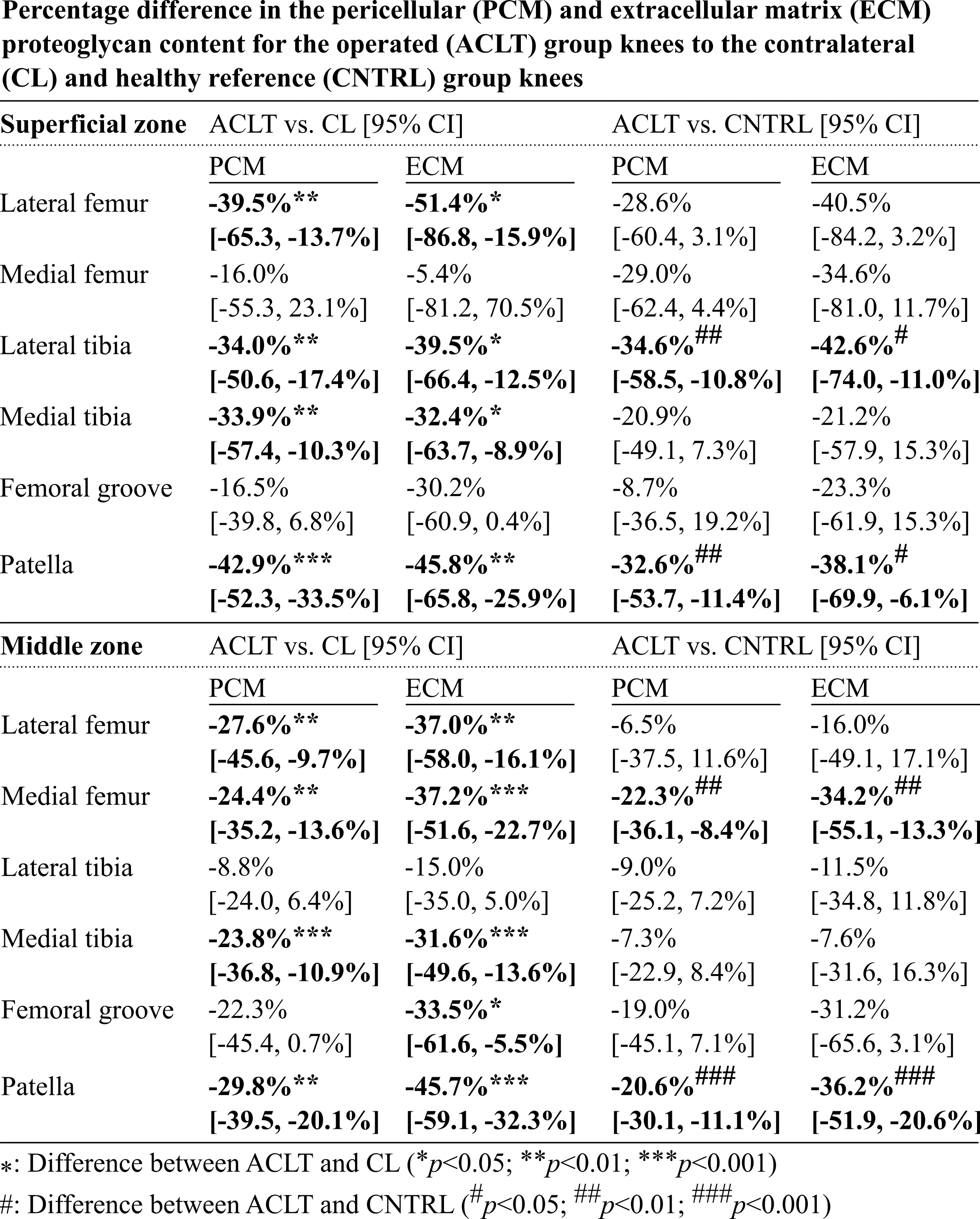

***Supplementary Table 4.*** *The mean percentage difference in pericellular (PCM) and extracellular (ECM) proteoglycan content with 95% confidence intervals of the anterior cruciate ligament transection operated (ACLT) to the contralateral (CL) and healthy control (CNTRL) group knees in superficial and middle zone cartilage for the lateral and medial femoral condyles, lateral and medial tibial plateaus, femoral groove, and patella.*

**B.2 Relative proteoglycan content of the cell microenvironment**

In the superficial zone, differences in the relative profiles between the ACLT and CNTRL groups were observed only near the cell edge or in the (presumed) territorial ECM (Supplementary Figure 3A). In the lateral femoral condyle, the relative proteoglycan content was higher in the ACLT group compared to the CL group (0.0–2.1 µm from the cell edge). Contrary to the lateral side, in the medial femoral condyle the CL group had higher relative proteoglycan content than the ACLT group (2.1–3.0 µm from the cell edge) and CNTRL group (0.9–3.0 µm from the cell edge).

In the middle zone cartilage, the relative proteoglycan content of the ACLT group knees was higher than in CNTRL group knees (Supplementary Figure 3B) for the medial femoral condyle (0.0–7.2 µm from the cell edge), the lateral femoral groove (0.0–4.3 µm from the cell edge), and the patella (0.0–5.5 µm from the cell edge). Additionally, relative proteoglycan content was higher in ACLT group knees than in CL group knees (up to 17.9 µm from the cell edge) at each knee location except at the lateral tibial plateau. The CL group knees showed smaller relative proteoglycan content compared to the CNTRL group knees near the cell edge for the lateral femoral condyle and further in the territorial matrix for the medial tibial plateau.

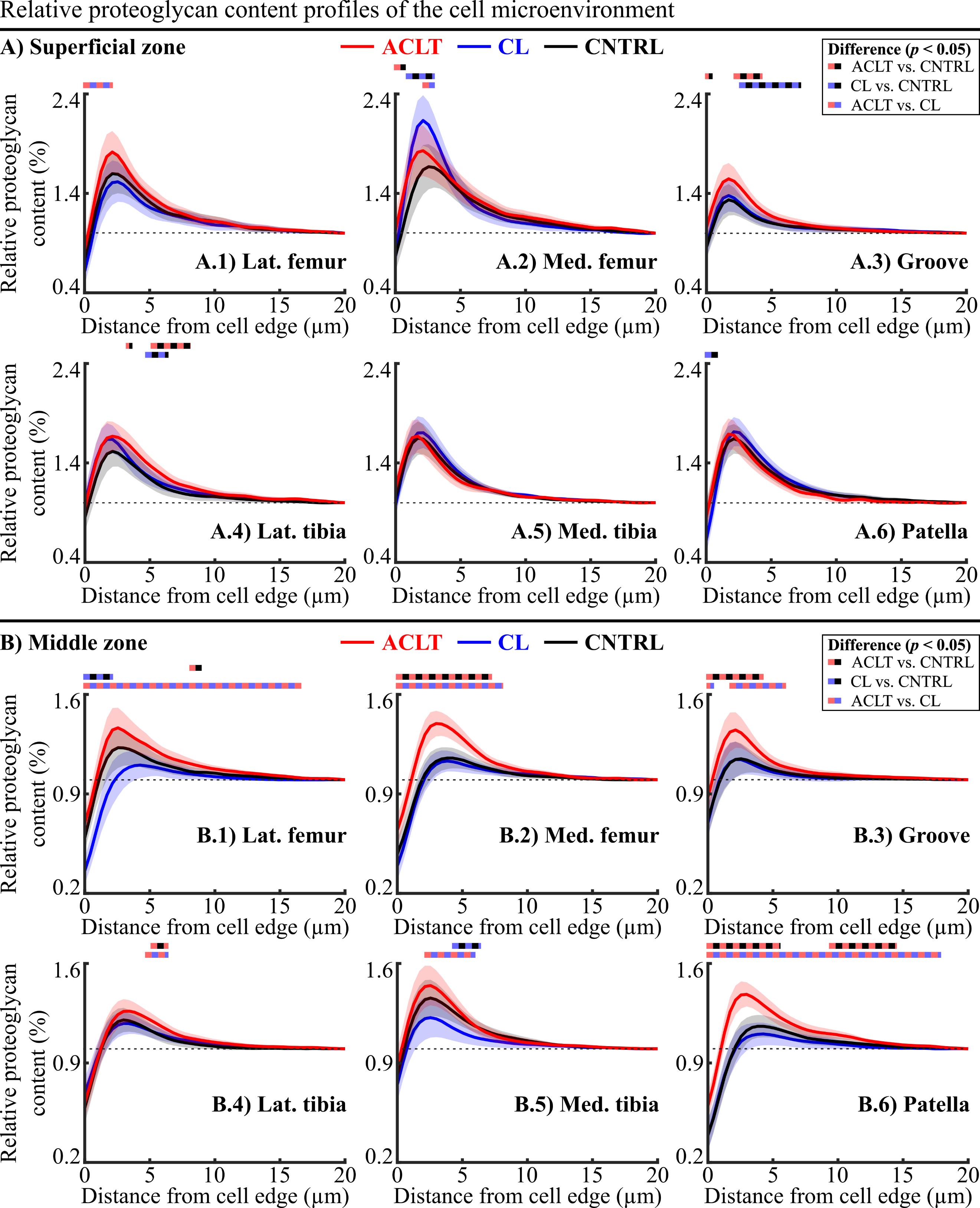

***Supplementary Figure 3.*** *Relative proteoglycan content profiles for the A) superficial and B) middle zone cell microenvironment in the 1) lateral femoral condyle, 2) medial femoral condyle, 3) lateral femoral groove, 4) lateral tibial plateau, 5) medial tibial plateau, and 6) patella. The mean profiles are drawn with solid lines for anterior cruciate ligament transection (ACLT, red), contralateral (CL, blue), and the separate healthy control (CNTRL, black) sample groups. Shaded areas of corresponding colors represent each mean profile's 95% confidence intervals. Dashed horizontal lines mark the point corresponding to the proteoglycan content of the extracellular matrix.*

**B.3. Associations between pericellular proteoglycan content and cell morphology in the superficial zone of cartilage**

Linear relationships between cell deformation and proteoglycan content of the cell microenvironment were mostly seen in the CNTRL and CL group knees at the different cartilage surfaces (Supplementary Figure 4). In the CNTRL group of the lateral femoral condyle, peak and ECM proteoglycan content were positively correlated with the size-related parameters of the chondrocytes (depth, area, and volume). In contrast, for the medial femoral condyle, a correlation was observed in the CNTRL group knees with more cell shape-related parameters like cell height, width, and aspect ratio. Interestingly, in the CNTRL group of the femoral groove, negative relationships were observed in volumetric and area change, which were positive in the lateral femoral condyle. Relationships were seldom observed in the tibial plateaus and patella.

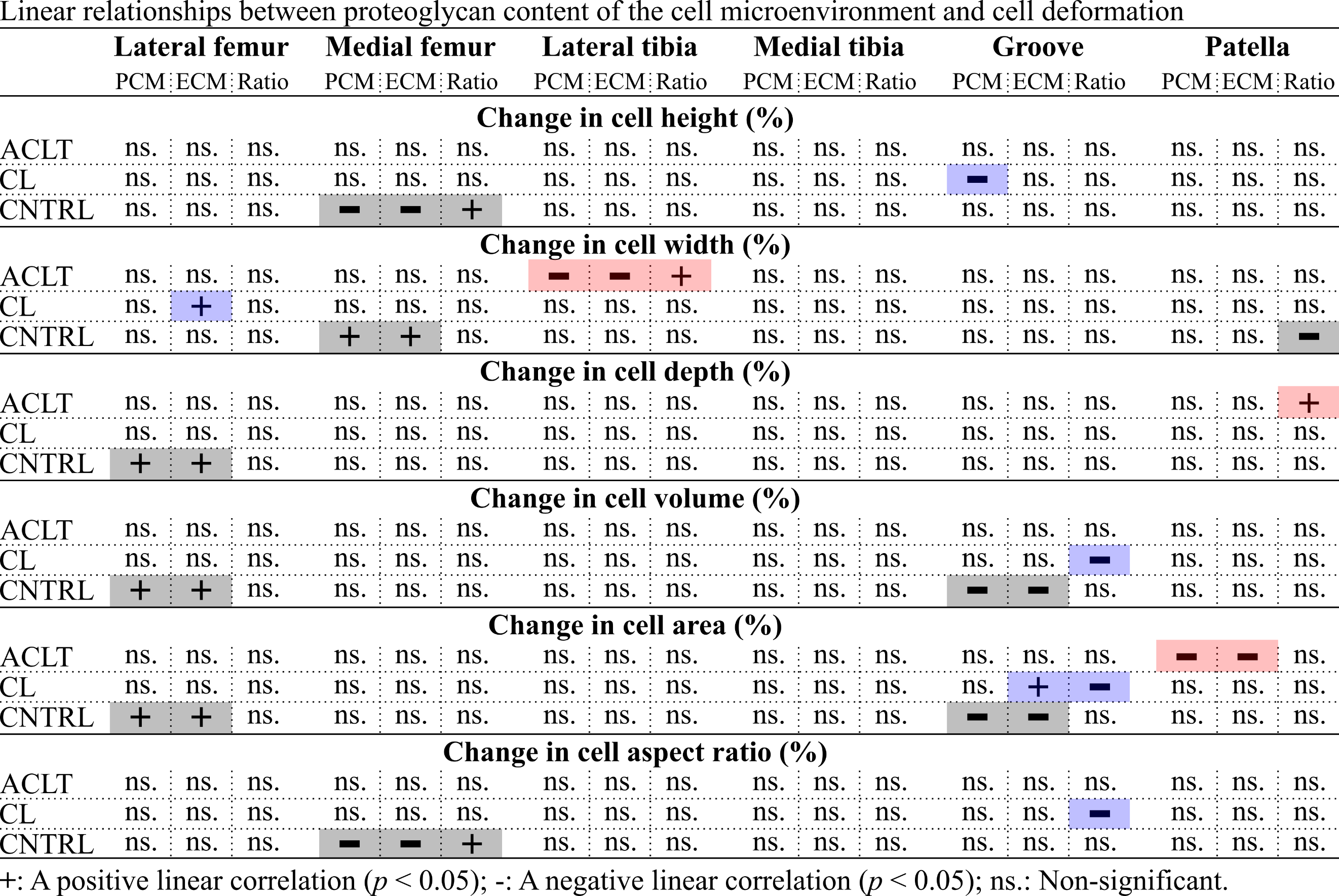

***Supplementary Figure 4.*** *Statistically significant associations (p < 0.05) observed between the proteoglycan content of the cell microenvironment and changes in analyzed cell morphology parameters in different joint cartilage sites with Pearson correlation analysis. Relationships are shown for the morphology parameters in each sample group (rows) at each cartilage surface (columns) for the peak proteoglycan content (PCM), proteoglycan content of the extracellular matrix (ECM) and the PCM/ECM ratio. Plus (+) and minus (-) signs indicate positive and negative linear correlations, respectively. Non-significant relationships are marked by ns.*

**References**

Han, S.-K., Colarusso, P., Herzog, W., 2009. Confocal microscopy indentation system for studying in situ chondrocyte mechanics. Medical Engineering & Physics 31, 1038–1042. https://doi.org/10.1016/j.medengphy.2009.05.013

Király, K., Lapveteläinen, T., Arokoski, J., Törrönen, K., Módis, L., Kiviranta, I., Helminen, H.J., 1996. Application of selected cationic dyes for the semiquantitative estimation of glycosaminoglycans in histological sections of articular cartilage by microspectrophotometry. The Histochemical Journal 28, 577–590. https://doi.org/10.1007/BF02331378

Kiviranta, I., Jurvelin, J., Säämänen, A.-M., Helminen, H.J., 1985. Microspectrophotometric quantitation of glycosaminoglycans in articular cartilage sections stained with Safranin O. Histochemistry 82, 249–255. https://doi.org/10.1007/BF00501401

Mäkelä, J.T.A., Rezaeian, Z.S., Mikkonen, S., Madden, R., Han, S.-K., Jurvelin, J.S., Herzog, W., Korhonen, R.K., 2014. Site-dependent changes in structure and function of lapine articular cartilage 4 weeks after anterior cruciate ligament transection. Osteoarthritis and Cartilage 22, 869–878. https://doi.org/10.1016/j.joca.2014.04.010

Ojanen, S.P., Finnilä, M.A.J., Mäkelä, J.T.A., Saarela, K., Happonen, E., Herzog, W., Saarakkala, S., Korhonen, R.K., 2020. Anterior cruciate ligament transection of rabbits alters composition, structure and biomechanics of articular cartilage and chondrocyte deformation 2 weeks post-surgery in a site-specific manner. Journal of Biomechanics 98, 109450. https://doi.org/10.1016/j.jbiomech.2019.109450

Ojanen, S.P., Finnilä, M.A.J., Reunamo, A.E., Ronkainen, A.P., Mikkonen, S., Herzog, W., Saarakkala, S., Korhonen, R.K., 2018. Site-specific glycosaminoglycan content is better maintained in the pericellular matrix than the extracellular matrix in early post-traumatic osteoarthritis. PLOS ONE 13, e0196203. https://doi.org/10.1371/journal.pone.0196203

Ronkainen, A.P., Fick, J.M., Herzog, W., Korhonen, R.K., 2016. Site-specific cell-tissue interactions in rabbit knee joint articular cartilage. Journal of Biomechanics 49, 2882–2890. https://doi.org/10.1016/j.jbiomech.2016.06.033
